# Zasp52’s differentially expressed intrinsically disordered region confers thin filament stability at the Z-disc

**DOI:** 10.64898/2026.03.10.710607

**Authors:** Nikolai Ho, Frieder Schöck

**Author notes:** Corresponding author: Frieder Schöck.

## Abstract

The Drosophila scaffolding protein Zasp52 is required to maintain structure at the muscle Z-disc which experiences strong forces during contraction. It is alternatively spliced into many isoforms, some of which contain a long intrinsically disordered region (IDR). We show that this region is primarily expressed in the indirect flight muscle (IFM) and is required for maintaining integrity of the Z-disc. Deleting the IDR-encoding exon 15e results in flightlessness and structural IFM defects, including sarcomere bending at the Z-disc and an inability to de-contract. These defects are indicative of a lack of proper thin filament anchoring to the Z-disc. This is further supported by a genetic interaction between exon 15e and actin. Fluorescence recovery after photobleaching of an isoform lacking exon 15e shows that the IDR is required for maintaining Zasp52 at the Z-disc and thereby stabilizing Z-discs. Lastly, we can rescue these phenotypes by restricting IFM use. Together, these results suggest that Zasp52’s IDR confers thin filament stability at the Z-disc of IFM.

## Background

The fruit fly *Drosophila melanogaster* possesses 152 muscle groups in the imago, as well as 54 in the larva (Bothe & Baylies, 2016). Each muscle is adapted to perform a specialized function, such as the extremely ordered indirect flight muscle (IFM) which is designed so as to enable the rapid oscillatory contractions needed to sustain flight. The marked phenotypic differences among these muscle types begs a molecular explanation. Alternative splicing is known to contribute to muscle type development and function with about 80% of sarcomere proteins being differentially expressed or spliced in Drosophila muscle (Nikonova et al., 2020). One such protein is Zasp52, a scaffolder, which contains numerous alternatively spliced exons generating a variety of isoforms.

All metazoan muscles share considerable structural homology (Steinmetz et al., 2012). They come in two types: striated, which appear striped given the presence of sarcomeres, and smooth, which do not possess sarcomeres. The sarcomere, a cylindrical structure, is the fundamental contractile unit of striated muscle and is bordered on either side by a dense circular structure called the Z-disc. Sarcomeres are arranged end-to-end in long arrays called myofibrils. Thin filaments composed of actin are anchored at and extend perpendicularly from the Z-disc into neighbouring sarcomeres, terminating at the H-zone (the region halfway between two Z-discs with no thin filaments). Thick filaments composed of muscle myosin are centred at the M-line (found at the centre of the H-zone) and interdigitate with thin filaments in a hexagonal lattice. The mechanism of sarcomere contraction is explained by the sliding filament theory in which thick filaments pull upon thin filaments in an ATP-dependent manner, bringing the Z-discs closer together. The simultaneous contraction of tandem sarcomeres comprising a myofibril is what leads to large-scale muscular contraction (Krans, 2010). The Z-disc is crucial for maintaining mechanical stability of the muscle and contains numerous proteins including Zasp52 which interacts with the thin filament cross-linker alpha-actinin (Liao et al., 2016) and actin (Liao et al., 2020) in Drosophila.

Zasp52, part of the three-membered Drosophila Alp/Enigma protein family along with Zasp66 and Zasp67, is crucial for maintaining structure at the Z-disc. It localizes to the Z-disc in all muscle types of *Drosophila melanogaster* (Katzemich et al., 2011) and is required for the assembly and maintenance of IFM sarcomeres (Katzemich et al., 2013; Liao et al., 2016; Medina-Quintana et al., 2026). Developmentally, Zasp52 appears in the embryo first at myotendinous junctions and also at Z-bodies - clusters which later coalesce to form Z-discs (Katzemich et al., 2013). Zasp52 binds actin via its N-terminal PDZ domain plus a long C-terminal extension (Liao et al., 2020), and α-actinin via a region encompassing the PDZ domain plus a short C-terminal extension (Liao et al., 2016). In addition to the PDZ and Zasp motif (ZM) domain, Zasp52 contains four LIM domains which interact with the ZM domain resulting in auto-oligomerization; this mechanism is believed to drive radial growth of the Z-disc as well as pathological aggregate formation (González-Morales et al., 2019; Medina-Quintana et al., 2026). Its primary human ortholog LDB3/ZASP is implicated in muscular dystrophies which are characterized by progressive adult-onset distal myofibrillar myopathy and sometimes cardiomyopathy (Griggs, et al., 2007).

Zasp52 is alternatively spliced into twenty-two different isoforms (according to FlyBase release FB2025_02, used throughout this paper), which leads to a variety of different domain and inter-domain combinations. The inter-domain linker regions connecting Zasp52’s structured domains (i.e., PDZ, ZM, and LIM domains) are less characterized but are proposed to play an important role in protein function (Fisher & Schöck, 2022).

Though not much is known about the linker region between LIM1 and LIM2, its predominant expression in IFM and unusually large length prompted us to investigate it. We were curious to see how deleting only this linker region while retaining all conserved domains would affect muscle structure and function. This linker was found to be highly disordered; intrinsically disordered regions (IDRs) have mostly been analyzed biochemically, with only limited data on their *in vivo* function in model organisms (Jensen et al., 2025). Flight ability was impaired and myofibrils displayed a variety of phenotypes which were all indicative of a loss of thin filament anchoring at the Z-disc. Most phenotypes exhibited an age-related increase in severity. Put together, these data demonstrate a crucial role for this large IDR in maintaining sarcomere integrity by stabilizing thin filaments at the Z-disc in IFM.

## Results

### exon 15e-containing isoforms are expressed in a specific spatiotemporal manner

Zasp52 isoforms containing the first and second LIM domains contain a linker region between the two which can vary greatly in size. Encoded by several exons, this linker region may contain exon 15 which undergoes alternative 5’ splice site selection generating five exonic variants (15a-e) ranging from 130 nucleotides (15a) to 4429 nucleotides (15e) in length **(Figure 1A)**. The last 29 nucleotides of exon 15 contribute to the second LIM domain, but the remainder encodes the linker. When exon 15e is included, it constitutes the majority of the entire protein, encoding 1475 amino acids which total 160 kDa. In the canonical and longest isoform Zasp52-PF, the entire protein is 2194 amino acids long and the linker region between LIM1 and LIM2 is 1686 amino acids long. Exon 15e is referred to variously by other sources: as exon 9 in Flybase, exon 16c in Katzemich et al., 2011, and exon 18/19 in Benna et al., 2009; protein isoforms containing exon 15e were previously also referred to as Z(210) (Chechenova et al., 2013).

**Figure 1.**
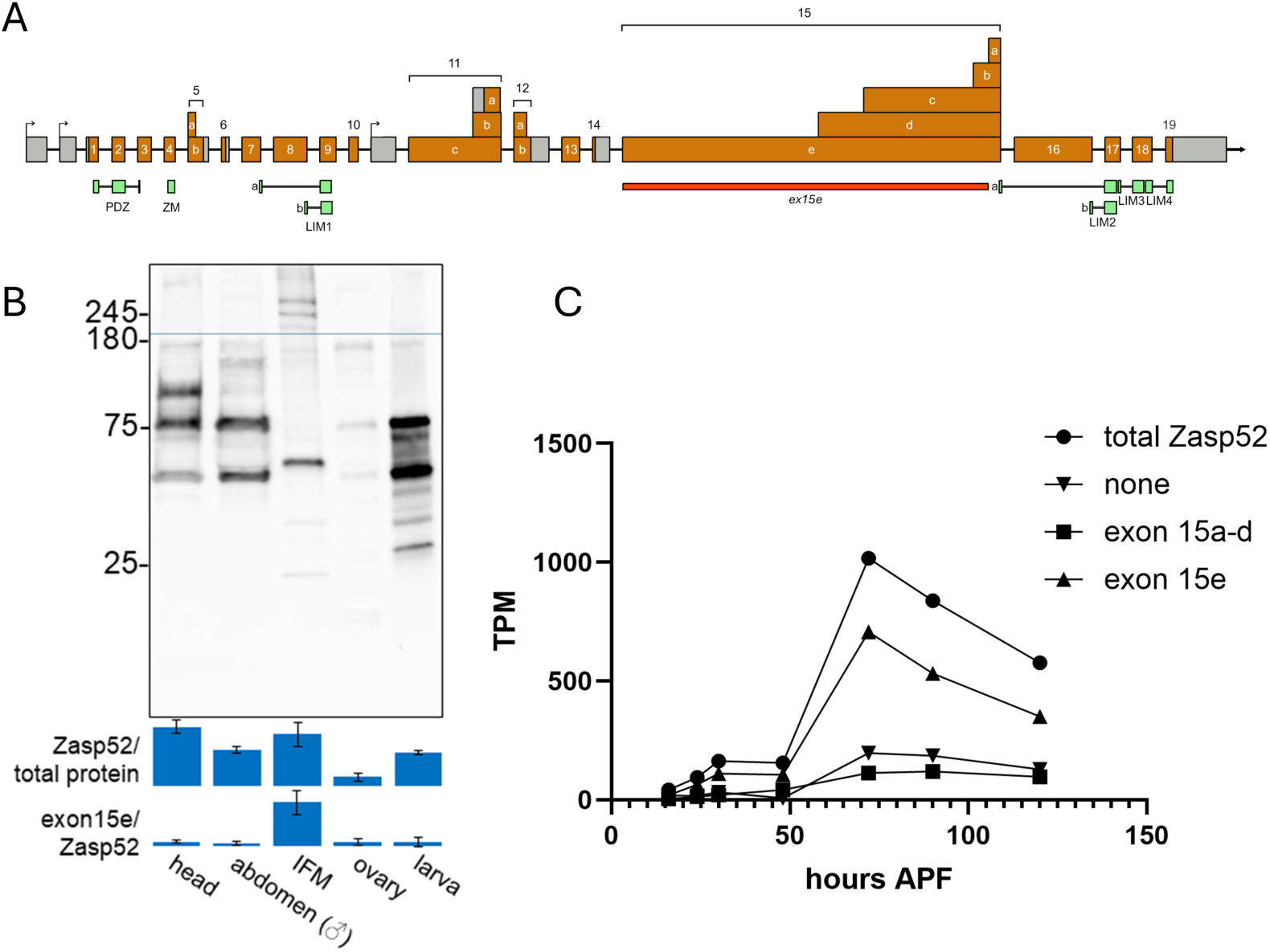
exon 15e-containing isoforms are expressed in a specific spatiotemporal manner. **(A)** A schematic of the *Zasp52* locus. Exons are drawn to scale, introns are not. Coding exons are shown in orange and untranslated regions in grey. Alternative start sites are also indicated. Below in green are mappings of structured domains according to SMART (Letunic & Bork, 2018); and also Katzemich et al, 2011 which defined LIM1 and LIM2 variants. The deletion in *ex15e* mutants is indicated in red. **(B)** Western blot using five different tissue extracts from control flies probed with a Zasp52 full-length antibody. Relative amounts of Zasp52 content as measured in each lane, over the total protein content as measured using the TGX Stain-free gel method, are quantified below for each tissue as mean with error bars representing standard deviation (*n*=3). Exon 15e content as measured using the protein content above the dotted blue line for each lane, over Zasp52 content is quantified below that. Molecular weights appear higher, perhaps due to retardation caused by non-specific interactions with the gel matrix, as also seen in Watts et al., 2017, Figure S1. **(C)** RNA-seq data from Spletter et al., 2018 of IFM tissue across various timepoints after puparium formation; the last timepoint is of adults one day after eclosion. Gene expression levels in transcripts per million (TPM) are shown for Zasp52 isoforms classified according to their splicing of exon 15. The majority of Zasp52 IFM isoforms contain exon 15e.

Western blotting using a polyclonal antibody capable of detecting all Zasp52 splice variants (hereinafter referred to as the Zasp52 full-length antibody, Jani & Schöck, 2007) was used to analyze the presence of Zasp52 in IFM, ovaries, whole larvae, heads, and whole male abdomens **(Figure 1B)**. IFM contains the second most Zasp52 protein of any isoform relative to total protein content compared to the other tissues analyzed and is the only tissue that significantly contains exon 15e-containing isoforms relative to total Zasp52 content. This agrees with previous studies (Chechenova et al., 2013, Benna et al., 2009) which found no higher weight isoforms in larvae, and with transcriptomics data. Whole organism RNA-seq data from the ENCODE project (Duff et al., 2015) reveals very little expression of exon 15e in all developmental stages up until four days past the white prepupal stage **(Figure 1 – Figure supplement 1)**. To verify that this is indeed a phenomenon of exon 15e inclusion and not merely Zasp52 expression, one can look at the surrounding exons which do not follow the same pattern as exon 15e. To gain a more precise perspective on IFM expression of exon 15e, we turned to an RNA-seq dataset from Spletter et al., 2018 which looked at total RNA content of IFM at various developmental time points **(Figure 1C)**. Expression of all Zasp52 transcripts in IFM surges from 48 to 72 h after puparium formation (APF), with most of these transcripts containing exon 15e, as opposed to including exon 15 a-d only or no exon 15 at all.

We used various software to predict the level of disorder across Zasp52-PF **(Figure 1 – Figure supplement 2A)**. All identified the linker region between LIM1 and LIM2 as an intrinsically disordered region (IDR) to various degrees, although some regions within it appear more ordered. The actin-binding motif (ABM) identified by Ashour et al., 2023 within exon 15e is predicted to be more structured than the surrounding regions. No significant sequence similarity was found when aligning Zasp52’s linker region to human or other *Drosophila melanogaster* ALP/Enigma protein family members. We then compared the lengths of the linker regions between LIM1 and LIM2 in all insect species possessing a Zasp52 ortholog as defined by NCBI’s Eukaryotic Genome Annotation Pipeline **(Figure 1 – Figure supplement 2B)**. Drosophilids generally have a longer linker than their taxonomical neighbours. Mayflies, dragonflies, and damselflies, which use direct flight muscle, have notably short linkers. Since linear sequence often does not fully describe an IDR’s function, we analyzed linear clustering or mixing of different residue types using NARDINI (Cohan et al., 2022). The z-score matrix of the linker region between LIM1 and LIM2 in Zasp52-PF showed a strong clustering/segregation of polar residues (µ), as well as alanine (A) and proline (P), and hydrophobic residues (h) **(Figure 1 – Figure supplement 2C)**. The z-score matrix of exon 15e alone displayed much of the same clustering, except with additional strong clustering of acidic residues (-) with respect to other residue classes **(Figure 1 – Figure supplement 2C’)**. This implies that the additional exons in the linker region dilute or mask the strong clustering of acidic residues in exon 15e alone. We attempted to compare z-score matrices across various arthropod species but were unable to detect any meaningful correlation indicating that the binary patterns uncovered by NARDINI do not reveal taxonomical differences in exon 15e composition.

### A CRISPR deletion of exon 15e displays flight defects

To further investigate the function of exon 15e, we generated a CRISPR deletion spanning five intronic nucleotides five prime to exon 15e up to 149 nucleotides before the 3’ end of exon 15e **(Figure 1A)**. In total, 4285 nucleotides totaling 156 kDa were deleted. Thus, our mutants would affect all isoforms containing exons 15b-e. The deletion endpoints were sequenced to confirm effective CRISPR knockout. To confirm that our deletion worked and spliced as intended, we performed Western blots using the Zasp52 full-length antibody to compare our deletion with a control and the MiMIC line *MI00979* (Venken et al., 2011) which truncates everything C-terminal to and including exon 15e **(Figure 2A)**. The largest protein that could possibly be present in our mutant would be 136 kDa (Zasp52-PI). Our deletion was devoid of all higher molecular weight isoforms containing exon 15e which were present in the control as expected. Our deletion also contained several isoforms which were heavier than the highest molecular weight isoforms of *MI00979* which were also not present in control, suggesting that these are novel products containing the C-terminal LIM domains. To confirm that this was indeed the case, i.e., that the three C-terminal LIM domains were retained, we raised an antibody against them and performed a Western blot **(Figure 2A’)**. Our CRISPR deletion retained these LIM domains as expected. We refer to this CRISPR mutant as *ex15e*.

**Figure 2.**
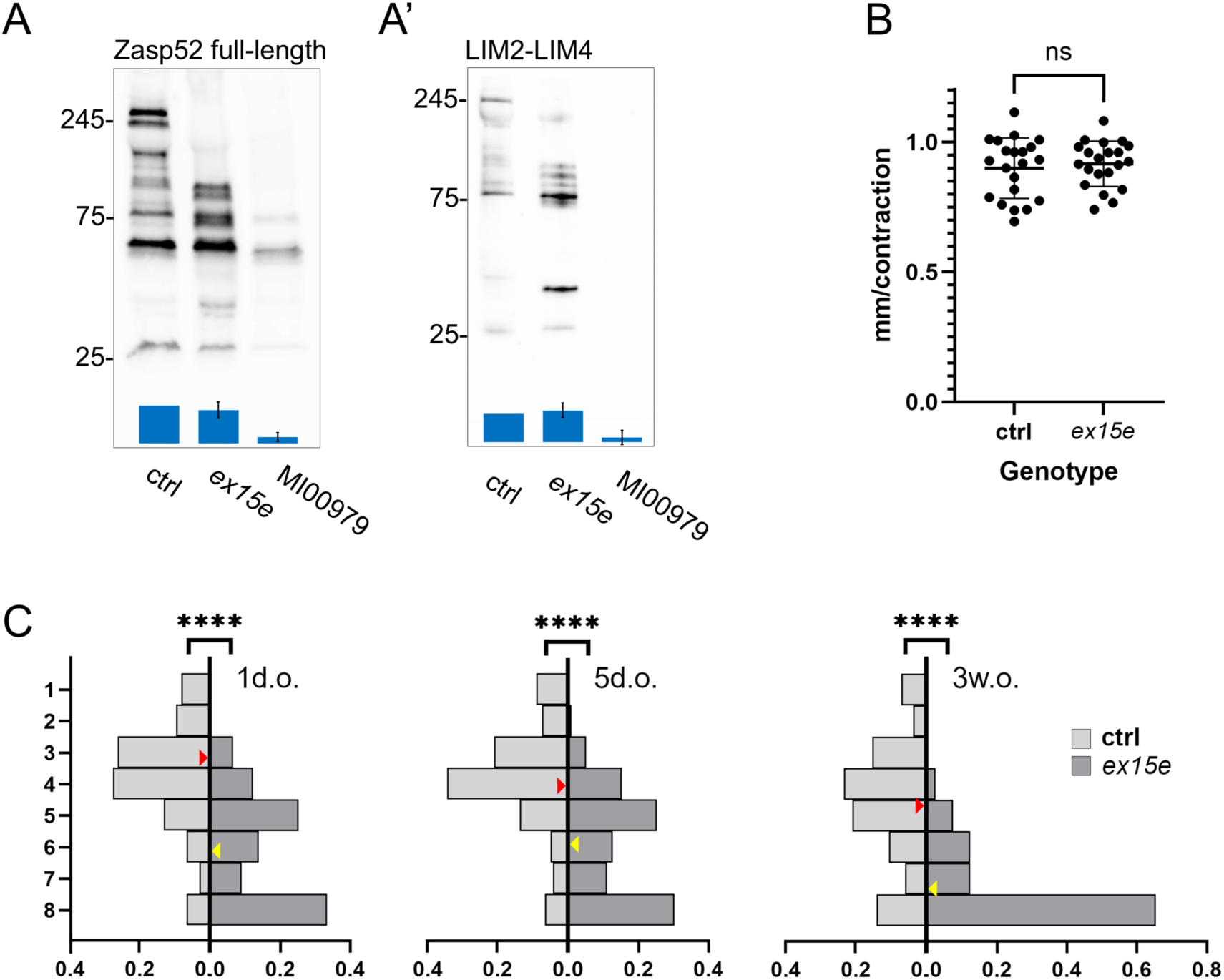
A CRISPR deletion of exon 15e displays flight defects. Western blots using either the Zasp52 full-length antibody **(A)** or antibody against LIM2-LIM4 **(A’)** of either *w^1118^* (ctrl hereinafter), *ex15e*, or the MiMIC line *MI00979* whole flies. Relative amounts of Zasp52 content as measured in each lane, over the total protein content, are quantified below for each genotype as mean with error bars representing standard deviation (*n*=3) normalized against the ctrl lane. **(B)** Larval crawling assay of either ctrl (*n*=21) or *ex15e* (*n*=21) third instar larvae showing no significant difference between the two (unpaired t-test with Welch’s correction; *p*=0.6006). **(C)** Flight assays of ctrl vs *ex15e* flies of three different age categories: one-day-old (ctrl *n*=138 flies, *ex15e n*=123), five-days-old (ctrl *n*=125, *ex15e n*=119), and three-weeks-old (ctrl *n*=86, *ex15e n*=81). The y-axis indicates flight strength: flies were released into a tube and those that landed in the top segment (y=1) had the strongest flight strength while those that landed in the bottom dish (y=8) had the weakest. The x-axis indicates the proportion of flies that landed in that segment. Red arrows indicate the average position landed in ctrl flies, yellow arrows are for *ex15e* flies. Difference in flight ability was statistically significant for all age categories (Fisher’s exact test; *p*<0.0001).

We wanted to see how *ex15e* affected the locomotory abilities of the fly. Since larvae do not express exon 15e, we do not expect them to be affected by our deletion. Therefore, we performed a larval crawling assay as a negative control to confirm that *ex15e* larval muscles were unaffected **(Figure 2B)**. As predicted, control and *ex15e* larvae demonstrated no differences in crawling ability. Next, we wanted to see how flight would be affected, given that exon 15e is present primarily in the adult IFM. *ex15e* flies of all age categories had impaired flight ability compared to control with a severe deterioration at three weeks of age **(Figure 2C)**. To ensure that no secondary-site mutations were affecting our mutant phenotypes, we performed a simple binary flight assay using the heterozygous *ex15e* deletion over a Zasp52 deficiency **(Figure 2 – Figure supplement 1A)**.

This impairment in flight could stem from a variety of causes, given the multimodal functionality of Zasp52. We decided to first look at gross muscle morphology before zooming in on our investigation. Thorax bisections revealed no noticeable difference in the IFMs of *ex15e* and control flies **(Figure 2 – Figure supplement 1B, C)**. Since Zasp52 was known to be involved in integrin interactions (Jani & Schöck, 2007; Bouaouina et al., 2012) and junctional components (Ashour et al., 2023), we inspected the myotendinous junctions of IFM, which are called modified terminal Z-discs (MTZ), in three-week-old flies **(Figure 2 – Figure supplement 1D-G’’)**. βPS integrin localization appeared unaffected in *ex15e*; this agrees with integrins being upstream of Zasp52 (Jani & Schöck, 2007). However, the MTZ appeared disrupted and malformed, lacking the organized structure seen in controls. Furthermore, we also noticed myofibril defects which prompted us to further investigate IFM at the myofibrillar level.

### *ex15e* mutant IFM phenotypes: bending at the Z-disc and thin filament intrusion into the H-zone

IFM tissue was analyzed by confocal microscopy with several phenotypes being observed. First, myofibrils in *ex15e* mutants were often found to be bent at the Z-disc **(Figure 3A-C)**. In control flies, sarcomeres across different age groups typically displayed the same proportion of degree of bentness. In *ex15e*, the proportion of bent sarcomeres increased with age, as well as the degree of bending, compared to control flies. Force-induced bending results in a sarcomere whose inner curve is bowed, much like a hose with a kink. The bending in *ex15e* appears to be a result of the improper anchoring of thin filaments which leads to a shear type angle shift in the sarcomere at the Z-disc. Bending was found exclusively at the Z-disc and not the M-line in all muscle analyzed.

**Figure 3.**
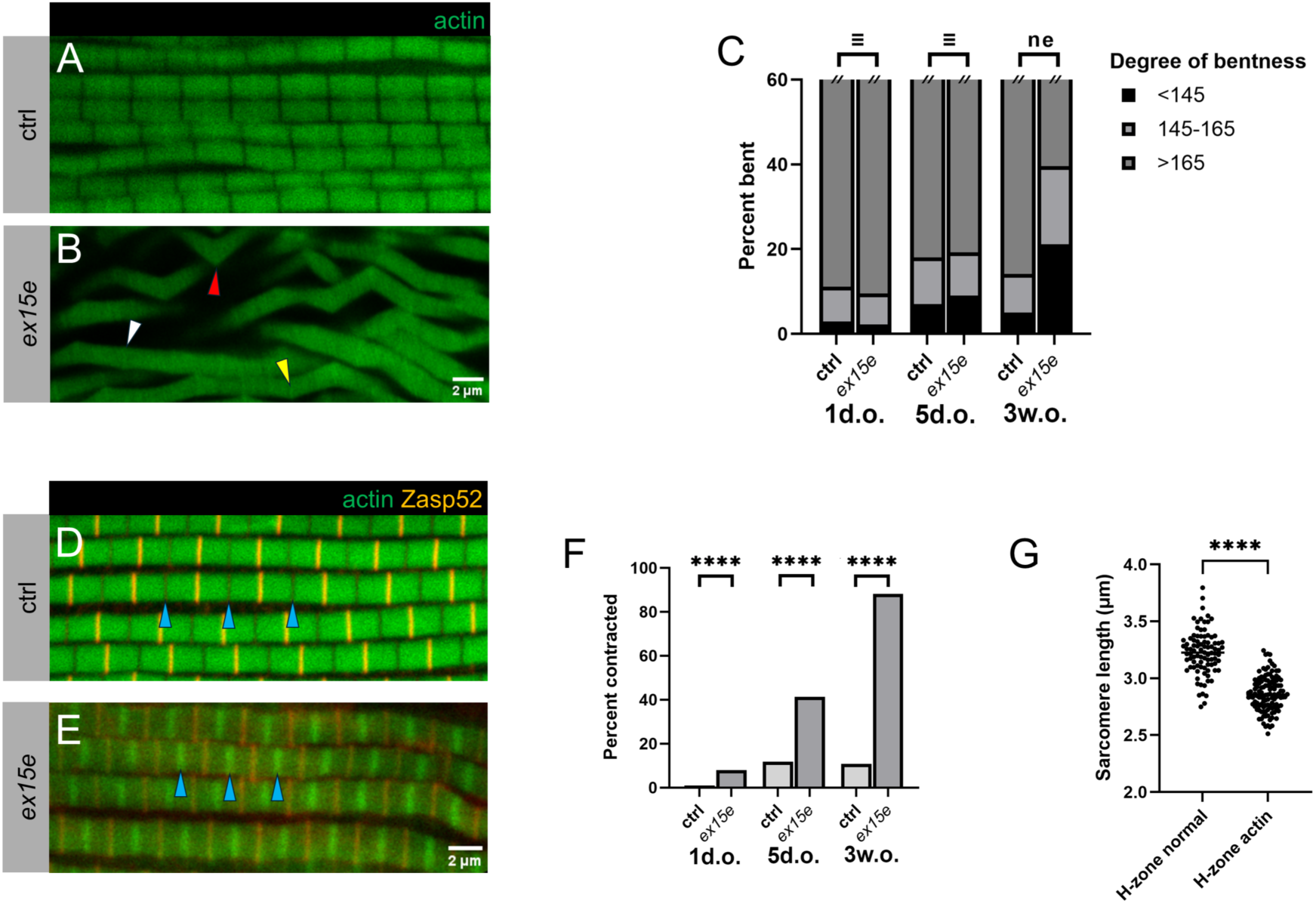
*ex15e* mutant IFM phenotypes: bending at the Z-disc and thin filament intrusion into the H-zone. **(A-C)** Analysis of bent sarcomere phenotype. Actin is visualized by phalloidin in green. Ctrl myofibrils **(A)** appear straight while *ex15e* myofibrils **(B)** contain a range of bending (representative three-week-old myofibrils shown). Red arrowhead indicates a representative sarcomere with a bend of <145°, yellow arrowhead indicates 145-165°, and white arrowhead >165°. **(C)** Bend angles were statistically equivalent (≡) between ctrl and *ex15e* in one-day-olds and five-day-olds, but not equivalent (ne) in three-week-olds, where *ex15e* displayed greater bentness. Sample sizes: one-day-old (ctrl *n*=1242 sarcomeres, *ex15e n*=1851), five-days-old (ctrl *n*=1192, *ex15e n*=946), and three-weeks-old (ctrl *n*=1768, *ex15e n*=1125) (Two One-Sided Test (TOST) with ±4.5° equivalence margin, α=0.05; 1d.o. *t_L_=*12.95, *t_U_*=-12.07; 5d.o. *t_L_=*6.03, *t_U_*=-8.01; 3w.o. *t_L_=*-6.34, *t_U_*=-20.63). **(D-F)** Sarcomeres also exhibited actin in the H-zone phenotype (representative three-week-old myofibrils shown). Ctrl myofibrils **(D)** display normal H-zones without actin with M-lines/H-zones indicated by blue arrowheads. *ex15e* myofibrils **(E)** display actin accumulation at the H-zone; arrowheads indicate same as above. Actin is visualized by phalloidin in green and Z-discs are identified by the full-length Zasp52 antibody in red. **(F)** These sarcomeres with H-zone actin, which appear hypercontracted, are significantly overrepresented in *ex15e* compared to ctrl across all age categories. Sample sizes were as follows: one-day-old (ctrl *n*=3255 sarcomeres, *ex15e n*=4056), five-days-old (ctrl *n*=3221, *ex15e n*=2677), and three-weeks-old (ctrl *n*=5394, *ex15e n*=4720 (Fisher’s exact test; *p*<0.0001). (**G)** To ascertain that sarcomeres with actin in the H-zone were indeed contracted, the Z-to Z-disc length was measured in *ex15e* three-week-old flies of both sarcomeres with a bare H-zone (H-zone normal, *n*=89 sarcomeres) and those with actin in the H-zone (H-zone actin, *n*=117). H-zone actin sarcomeres were significantly smaller (unpaired t-test with Welch’s correction; *p*<0.0001).

Another phenotype observed in *ex15e* was the presence of a dense band of actin staining at the H-zone, where normally there would be no actin staining **(Figure 3D-F)**. Control flies have very few sarcomeres with this phenotype at all ages; *ex15e* are similar to control at one day of age but begin to accumulate a large number of these abnormal sarcomeres with time. At three weeks of age, a large proportion of sarcomeres have actin at the H-zone. Usually, groups of myofibrils in the same area would exhibit this phenotype rather than these abnormal myofibrils being randomly dispersed among the normal ones. This dense band of actin suggested that thin filaments were overlapping at the M-line. This could be a result of hypercontraction, where thin filaments are brought so close together that they overlap. However, the protocol used for these preparations includes a relaxing solution containing ATP, which induces cross-bridge detachment resulting in the adoption of a relaxed conformation. To see if these abnormal sarcomeres with actin in the H-zone were indeed contracted, we measured their length and compared them to normal sarcomeres from the same sample in three-week-old flies **(Figure 3G)**. Sarcomeres with actin in the H-zone had a smaller length than those without indicating that they were indeed contracted. Muscle decontraction relies on the elastic nature of the sarcomere to restore it to a relaxed conformation once myosin heads detach from the thin filaments. This requires stable Z-discs and myofilament attachments, the lack of which may be the cause of a permanent hypercontracted state in *ex15e*.

To further investigate these phenomena, we performed transmission electron microscopy (TEM) of myofibrils and noticed phenotypes that concurred with our immunofluorescence analysis. Wild type IFM sarcomeres of three-week-old flies are highly stereotypic with electron-dense, continuous Z-discs and regular H-zones **(Figure 4A)**. In contrast, *ex15e* mutants show low-density, discontinuous Z-discs with many gaps as well as irregular H-zones **(Figure 4B)**. We also observed bent Z-discs and split myofibrils, as well as fraying of filaments at the sarcomere periphery **(Figure 4C)**. The latter two phenotypes could also be observed in transverse sections of myofibrils **(Figure 4D)**. Altogether, these observations indicate a breakdown of Z-disc coherence and stability leading to additional downstream effects like myofibril splitting and fraying.

**Figure 4.**
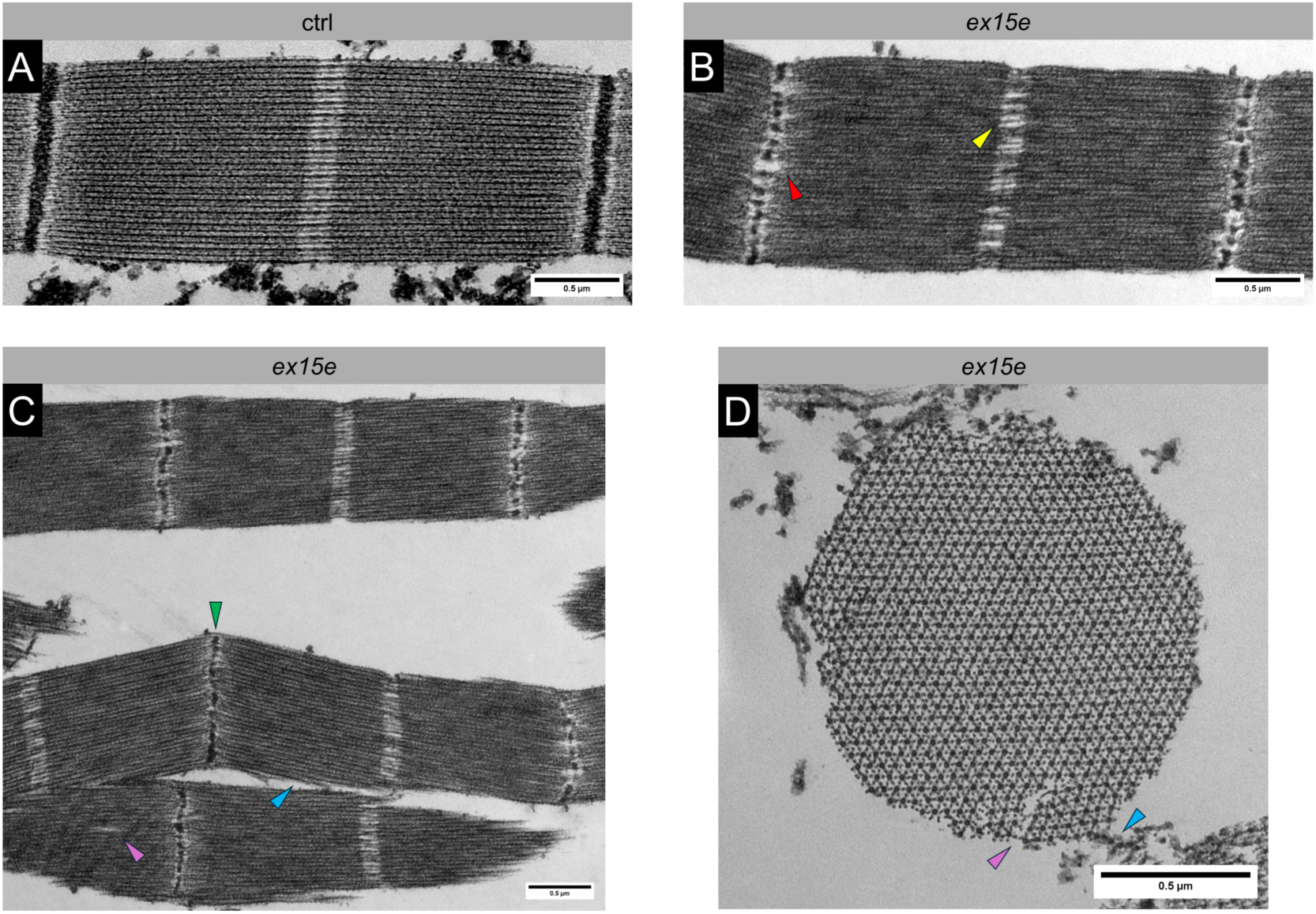
Transmission electron microscopy reveals defects in mutant sarcomere ultrastructure. TEM images reveal similar phenotypes as immunofluorescent ones (three-week-old shown). Ctrl **(A)** sarcomeres are very structured and even. Longitudinal sections of *ex15e* **(B, C)** display broken Z-discs (red arrowhead), broken M-lines (yellow arrowhead), bending at the Z-discs (green arrowhead), splits in filaments (purple arrowhead), and fraying of peripheral filaments (blue arrowhead). Transverse sections of *ex15e* **(D)** also display some of these abnormalities (purple and blue arrowhead).

### A putative actin-binding motif within exon 15e is not responsible for *ex15e* defects

A putative F-actin binding motif (ABM) identified by Ashour et al., 2023 is located within exon 15e and might be responsible for the thin filament-related phenotypes we witnessed in *ex15e* (see **Figure 1- Figure supplement 2A** for location). This short motif found in the CapZbeta protein of various metazoans is present in the Zasp52 proteins of the *D. melanogaster* subgroup (Ashour et al., 2023). To test the role of this ABM, we used CRISPR to delete five amino acids (LQREL) which include two highly conserved leucines and confirmed the in-frame deletion by sequencing – we refer to this mutant as *ΔABM*. Myofibrils appear normal **(Figure 5A-C)** and do not display the abnormal bending **(Figure 5D)** and contraction **(Figure 5E)** of *ex15e*. Flight ability distributions are slightly different from control flies although the average flight index is almost identical **(Figure 5F)**. In conclusion, *ΔABM* does not phenocopy *ex15e* which indicates that exon 15e’s function is not reducible to that short motif. Instead, the IDR is essential for the phenotypes we observed.

**Figure 5.**
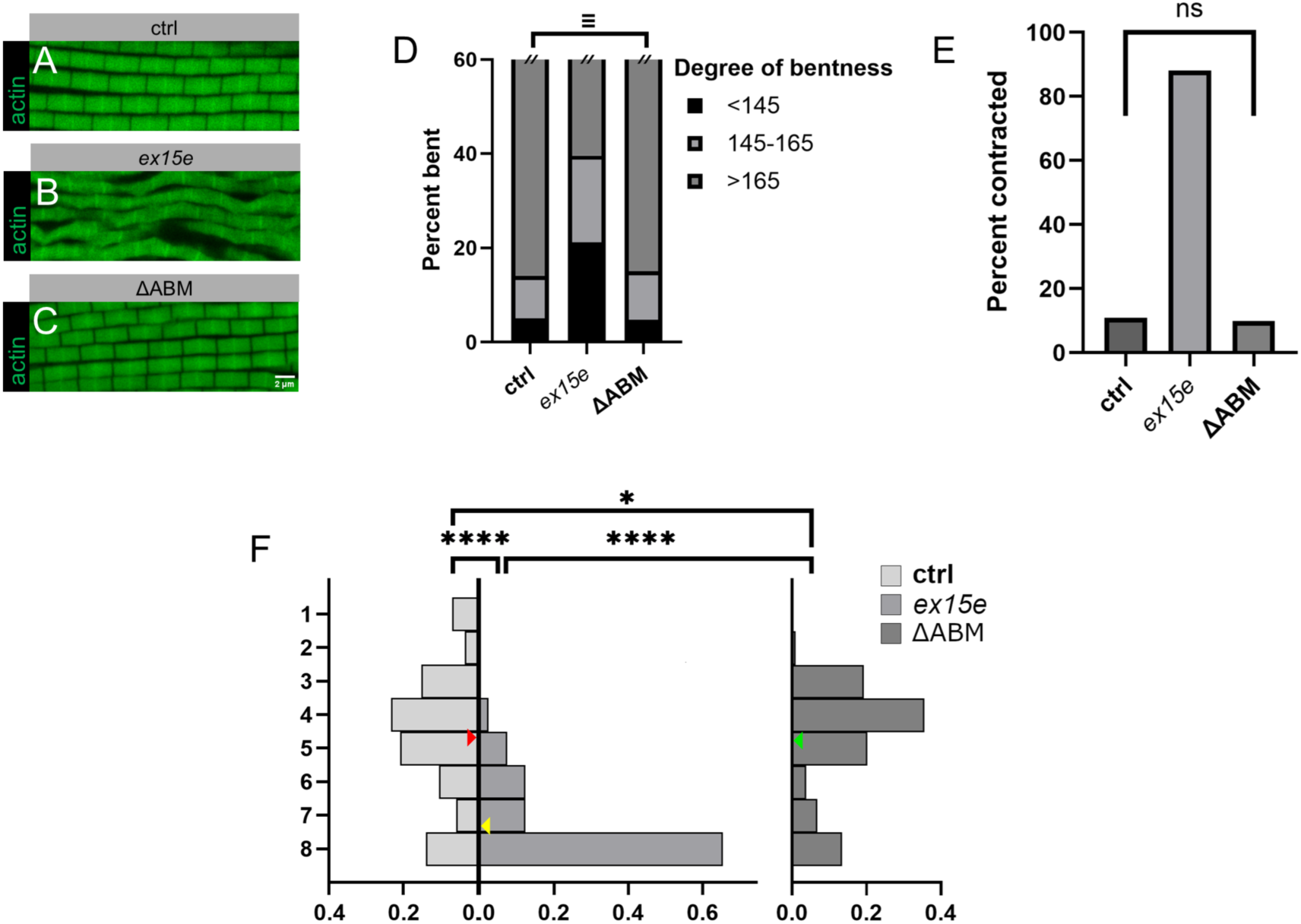
A putative actin-binding motif within exon 15e is not responsible for *ex15e* defects. *ΔABM* homozygotes along with all other flies in this figure were analyzed at three weeks of age. *ΔABM* myofibrils **(C)** resembled those of ctrl **(A)** and showed none of the phenotypes seen in *ex15e* mutants **(B)**. Actin is visualized by phalloidin in green. **(D)** Bend angles were equivalent between ctrl and *ΔABM* flies (*n*=1648) (Two One-Sided Test (TOST) with ±4.5° equivalence margin, α=0.05; *t_L_=*21.88, *t_U_*=-7.93). **(E)** The proportion of sarcomeres that were contracted was not significantly different between ctrl and *ΔABM* flies (*n*=3773) (Fisher’s exact test; *p*=0.1274). **(F)** Flight ability was only slightly decreased in ΔABM flies (*n*=104 flies) compared to ctrl (*n*=86) but significantly improved compared to *ex15e* (*n*=81). Red arrows indicate the average position landed in ctrl flies, yellow are *ex15e*, and green are *ΔABM* (Fisher’s exact test; ctrl-*ΔABM p*=0.0351; ctrl-*ex15e* and *ex15e*-*ΔABM p*<0.0001).

### *ex15e* defects are restored by rescue constructs

To determine if the phenotypes are truly due to the IDR or are caused by an altered isoform composition, we wanted to see if the phenotypes of *ex15e* could be rescued with two different isoforms: Zasp52-PF is the canonical full-length Zasp52 isoform which contains exon 15e, while Zasp52-PR contains all structured domains but not exon 15e **(Figure 6A)**. We expressed either UAS-Zasp52-PF or UAS-Zasp52-PR using the UH3-GAL4 driver in a homozygous *ex15e* background and examined three-week-old flies **(Figure 6B-H)**. UH3-GAL4 is expressed in IFM from 36 hours APF into adulthood (Singh et al., 2014). The degree of bending in Zasp52-PR rescue sarcomeres was statistically equivalent to that of *ex15e* mutants, while that of Zasp52-PF rescues was equivalent to control flies, indicating that Zasp52-PF but not Zasp52-PR can rescue this phenotype **(Figure 6F)**. The proportion of H-zone actin (i.e., hypercontracted) sarcomeres was not significantly different between *ex15e* and Zasp52-PR rescues as expected. Interestingly, no hypercontracted sarcomeres were observed upon re-expression of Zasp52-PF again indicating a successful rescue **(Figure 6G)**. Finally, we tested flight ability and observed a significant increase in flight strength in Zasp52-PF rescues compared to Zasp52-PR rescues, with the latter being not significantly different from *ex15e* **(Figure 6H)**. Zasp52-PF rescues were still weaker fliers than control flies suggesting a partial rescue of flight ability. Overall, only a transgene containing exon 15e resulted in rescue, showing that the phenotypes are due to the IDR.

**Figure 6.**
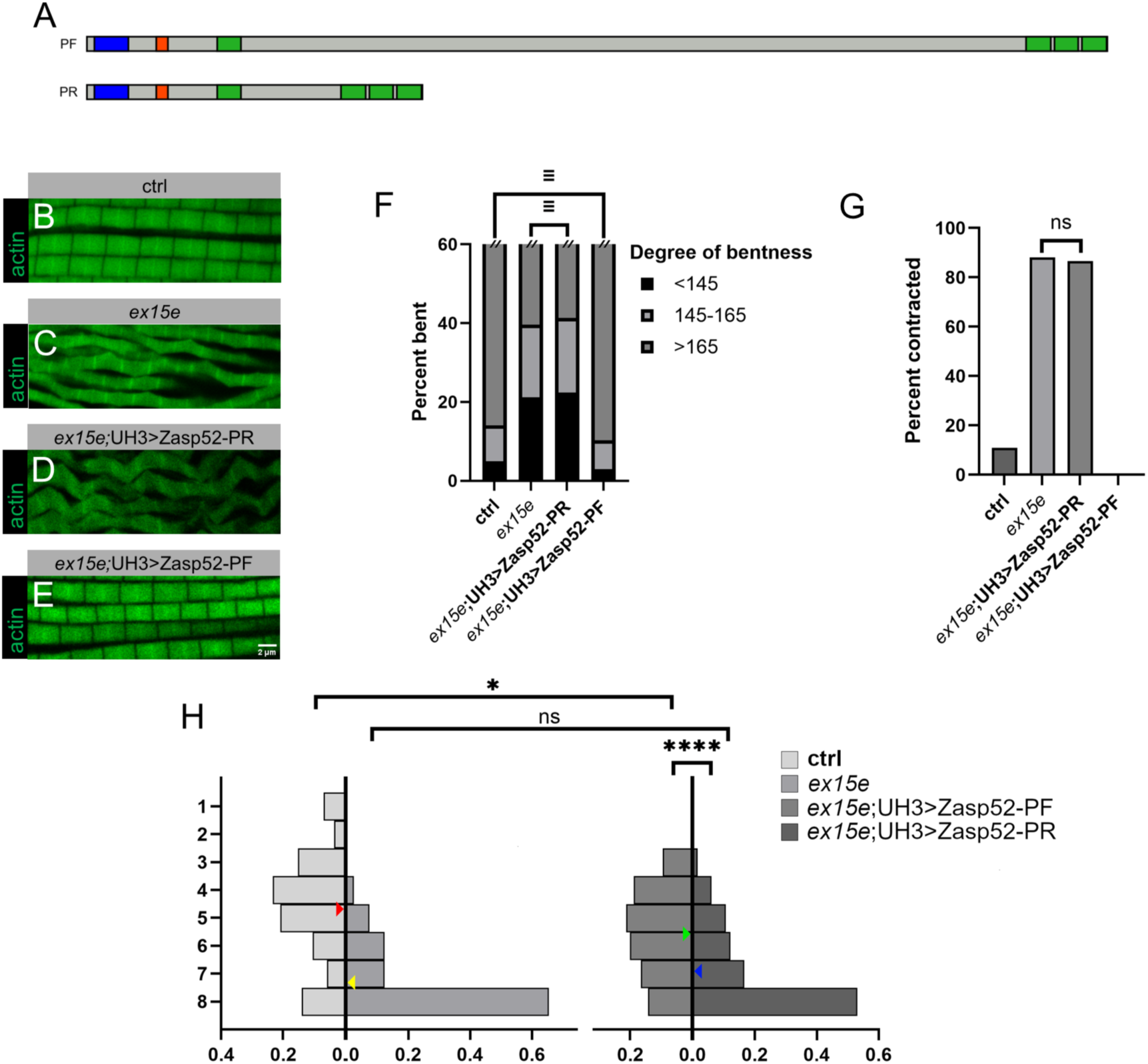
*ex15e* defects are restored by rescue constructs. Zasp52-PF or Zasp52-PR was heterozygously overexpressed with a UH3-GAL4 driver in a homozygous *ex15e* background in a rescue attempt. In this figure, all myofibrils had actin visualized with phalloidin in green. All experiments were done at three weeks of age, and ctrl and *ex15e* data are same as previous. **(A)** Schematic showing Zasp52-PF which is the canonical full-length isoform, and Zasp52-PR which contains all structured domains but lacks exon 15e. PDZ domain (blue), ZM (red), LIM domains (green). **(B-E)** Myofibrils display bending, Z-disc disruption, and overall myofibrillar disorganization in Zasp52-PR overexpressions **(D)**, similar to *ex15e* **(C)**, while Zasp52-PF overexpression **(E)** appears similar to ctrl **(B)**. **(F)** Bend angles were equivalent (≡) between *ex15e* and *ex15e*; UH3>Zasp52-PR, and between ctrl and *ex15e*; UH3>Zasp52-PF. Sample sizes: *ex15e*; UH3>Zasp52-PR *n*=2049 sarcomeres, *ex15e*; UH3>Zasp52-PF *n*=2221 (Two One-Sided Test (TOST) with ±4.5° equivalence margin, α=0.05; *ex15e*-*ex15e*; UH3>Zasp52-PR *t_L_=*1.92, *t_U_*=-10.16; ctrl-*ex15e*; UH3>Zasp52-PF *t_L_=*17.65, *t_U_*=-8.51). **(G)** The proportion of sarcomeres that were contracted (i.e., actin in H-zone) was not significantly different between *ex15e* and *ex15e*-*ex15e*; UH3>Zasp52-PR, whereas no such sarcomeres were found in *ex15e*; UH3>Zasp52-PF. Sample sizes: *ex15e*; UH3>Zasp52-PR *n*=2843 sarcomeres, *ex15e*; UH3>Zasp52-PF *n*=2480 (Fisher’s exact test; *p*=0.0521). **(H)** Flight ability was significantly improved in *ex15e*; UH3>Zasp52-PF (*n*=85) compared to *ex15e*; UH3>Zasp52-PR (*n*=66), with *ex15e*; UH3>Zasp52-PR showing no significant difference from *ex15e* but *ex15e*; UH3>Zasp52-PF showing significant difference from ctrl. Red arrows indicate the average position landed in ctrl flies, yellow are *ex15e*, green are *ex15e*; UH3>Zasp52-PR, and blue are *ex15e*; UH3>Zasp52-PF (Fisher’s exact test; *ex15e*; UH3>Zasp52-PR-*ex15e*; UH3>Zasp52-PF *p*<0.0001; *ex15e*-*ex15e*; UH3>Zasp52-PR *p*=0.2668; ctrl-*ex15e*; UH3>Zasp52-PF *p*=0.0129).

### *ex15e* and actin genetically interact

We next wanted to test for a genetic interaction between exon 15e and Act88F (the actin isoform present in IFM) by analyzing heterozygous *ex15e*, heterozygous *Act88F^KM88^* (an amorphic mutant of *Act88F*), and a di-heterozygous combination of both (*ex15e*/+; *Act88F^KM88^*/+) **(Figure 7A-D)**. *ex15e*/+ resembled control myofibrils and *Act88F^KM88^*/+ has mild myofibrillar defects. *ex15e*/+; *Act88F^KM88^*/+ exhibited supra-additive effects where we often noticed a novel phenotype of very large H-zones. A ratio of twice the thin filament length to the total sarcomere length was computed for all genotypes. Lower values represented sarcomeres whose thin filaments occupied a smaller proportion of the sarcomere and consequently had larger H-zones, whereas a value of one represented the absence of an H-zone. This ratio was not significantly different between control, *ex15e*/+, and *Act88F^KM88^*/+, but was considerably smaller in di-heterozygotes **(Figure 7E)**, indicating a strong genetic interaction between Act88F and exon 15e.

**Figure 7.**
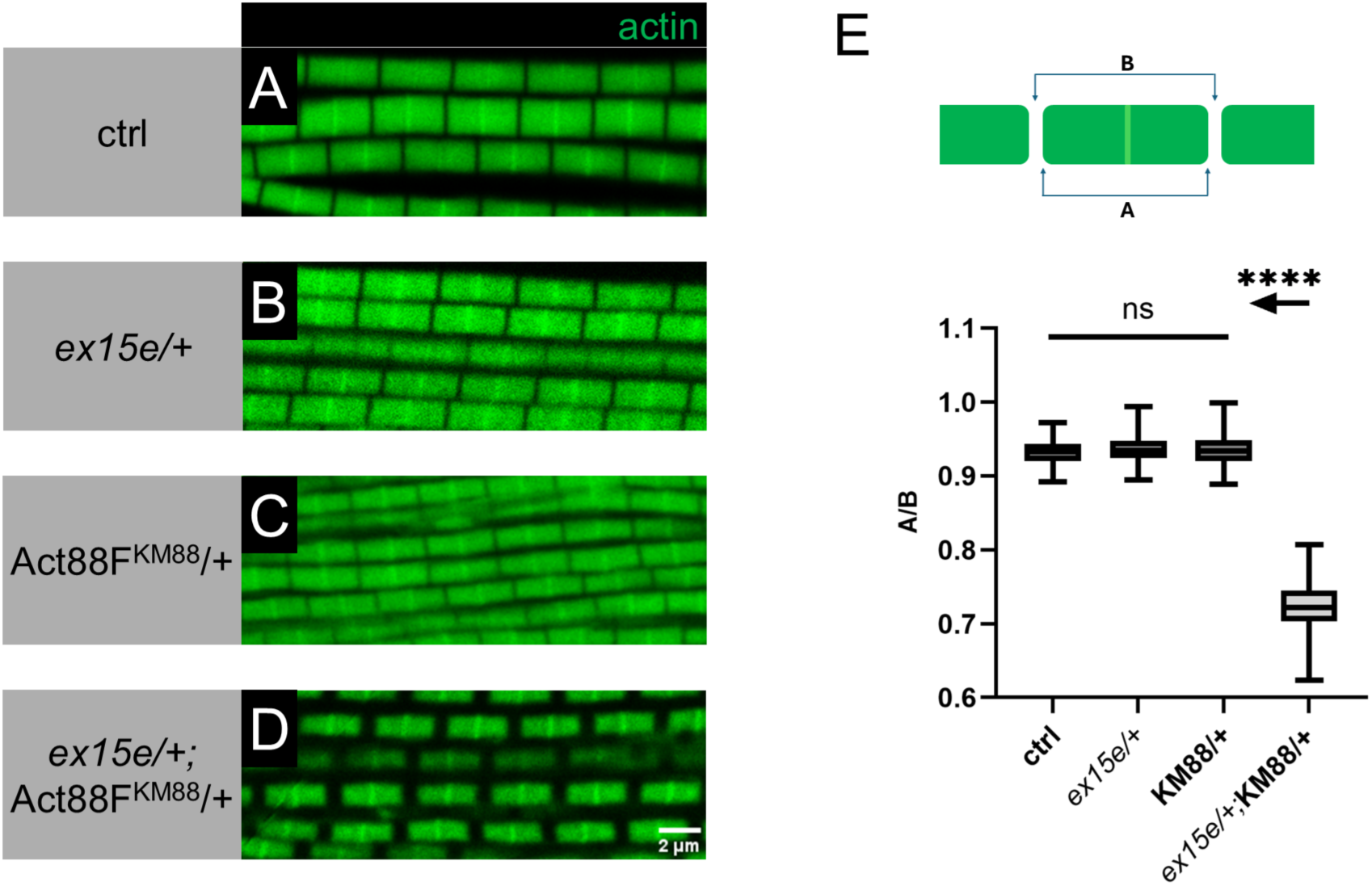
Large H-zones demonstrate a genetic interaction between exon 15e and Act88F. **(A-D)** Myofibrils of one-day-old flies of various genotypes. Actin was visualized with phalloidin in green. Ctrl **(A)** and heterozygous *ex15e* myofibrils **(B)** appear indifferentiable; heterozygous *Act88F^KM88^* **(C)** are slightly disrupted with fraying and slight narrowing. Representative *ex15e*/+; *Act88F^KM88^*/+ myofibrils **(D)** with large H-zones are shown. **(E)** We quantified the ratio of twice the thin filament length denoted distance “A” to the corresponding sarcomere’s length denoted as “B” for all four genotypes. A schematic illustrates the measurement of these distances. Sample sizes were as follows: ctrl *n*=119 sarcomeres, *ex15e*/+ *n*=122, *Act88F^KM88^*/+ *n*=119, *ex15e*/+; *Act88F^KM88^*/+ *n*=119. The ratio A/B represents the proportion of the sarcomere length occupied by thin filaments and is inversely correlated with H-zone length; *ex15e*/+; *Act88F^KM88^*/+ sarcomeres have significantly larger H-zones than all others (unpaired t-test with Welch’s correction; ctrl-*ex15e*/+ *p*=0.0502; ctrl-*Act88F^KM88^*/+ *p*=0.2770; ctrl-*ex15e*/+; *Act88F^KM88^*/+ *p*<0.0001; *ex15e*/+-*Act88F^KM88^ p*=0.4938; *ex15e*/+-*ex15e*/+; *Act88F^KM88^ p*<0.0001; *Act88F^KM88^*/+- *ex15e*/+; *Act88F^KM88^ p*<0.0001). All sarcomeres with an *Act88F^KM88^* allele were radially narrower than ctrl, but a synthetic enhancement could not be concluded from this observation.

### FRAP reveals defects in *ex15e* protein dynamics

We decided to test whether exon 15e could localize to the Z-disc on its own. To do this, we expressed the GFP-tagged exon 15e using the UH3-GAL4 driver (UH3>GFPexon15e). We observed successful localization to the IFM Z-discs **(Figure 8A)** and thus decided to assess kinetics using Fluorescence Recovery After Photobleaching (FRAP). One-day-old IFM Z-discs were photobleached and recovery of fluorescence was monitored. Compared to GFP-tagged Zasp52-PR (UH3>GFPZasp52-PR) which lacks exon 15e, exon15e-GFP displayed much lower recovery over a 96 second period **(Figure 8B)**. This suggests that the IDR stabilizes Zasp52 at the Z-disc and reduces its turnover. We then expressed GFP-tagged Zasp52-PR, which cannot rescue (see **Figure 6**), in an *ex15e* background (*ex15e*; UH3>GFPZasp52-PR). Fluorescence recovery in the *ex15e* background was to a significantly higher level compared to the wild type background **(Figure 8C)**, resulting in a larger mobile fraction **(Figure 8D)**. A greater mobile fraction in our mutant background confirms that the IDR is required to hold Zasp52 in place at the Z-disc, either by fastening itself, and/or keeping other molecules in place thereby contributing to overall Z-disc stability.

**Figure 8.**
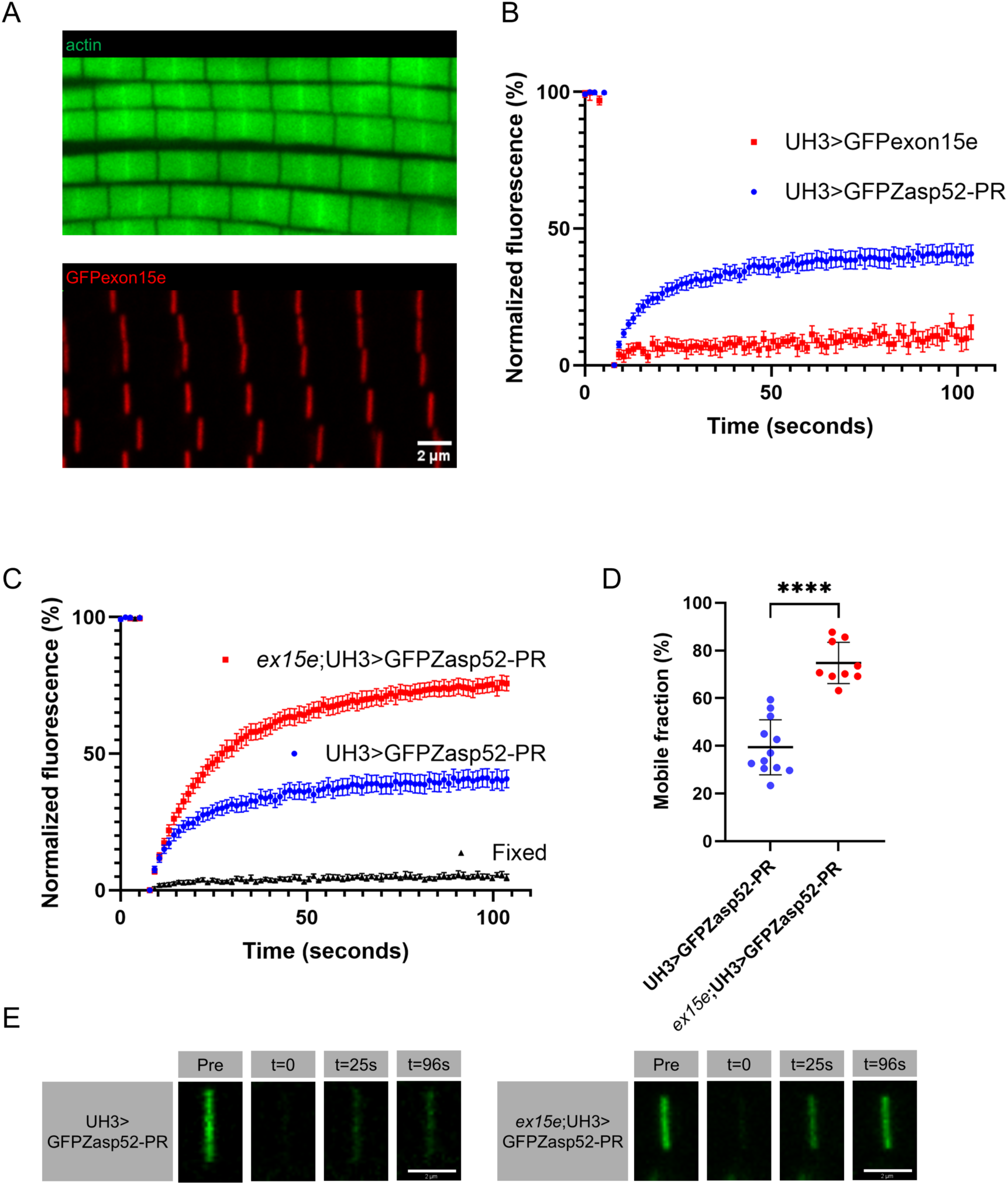
FRAP reveals defects in *ex15e* protein dynamics. **(A**) GFP-tagged exon 15e localized to the IFM Z-discs in one-day-old females. **(B)** FRAP experiments visualizing recovery of either GFP-tagged exon 15e (UH3>GFPexon15e, *n*=8) or GFPZasp52-PR (UH3>GFPZasp52-PR, *n*=12) were performed on one-day-old female IFM Z-discs. **(C)** Recovery of GFPZasp52-PR was also assessed in one-day-old female IFM Z-discs either in our mutant background (*ex15e*;UH3>GFPZasp52-PR, *n*=9 flies) or wild-type background (same data as in **B**). Ctrl UH3>GFPZasp52-PR Z-discs fixed with 4% paraformaldehyde for 10 minutes (labelled “Fixed”, *n*=7) displayed no recovery as expected. **(D)** Plateaus of fitted curves were significantly different, with a greater mobile fraction in the mutant background (unpaired t-test with Welch’s correction; *p*<0.0001). **(E)** Representative snapshots of bleached Z-disc recovery illustrate this difference between genotypes.

### *ex15e* defects are rescued by immobilization

On a different note, we noticed that virtually all the *ex15e* phenotypes worsened with age, which suggests that use might contribute to deterioration. We therefore decided to prevent flies from using their IFMs by immobilizing them. Flies were reared until three weeks of age in between glass plates separated by 1.5 mm spacers which prevented them from flying or even attempting to fly. Myofibrils appear normal in immobilized *ex15e* mutant flies **(Figure 9A-C)**. The degree of bending in immobilized flies was equivalent to that of control flies **(Figure 9D)**. The proportion of hypercontracted sarcomeres was also not significantly different between immobilized and control flies **(Figure 9E)**. Intriguingly, even flight ability was restored in immobilized flies compared to age-matched controls with no significant difference between the two **(Figure 9F)**. These results suggest successful rescue by preventing IFM use via immobilization.

**Figure 9.**
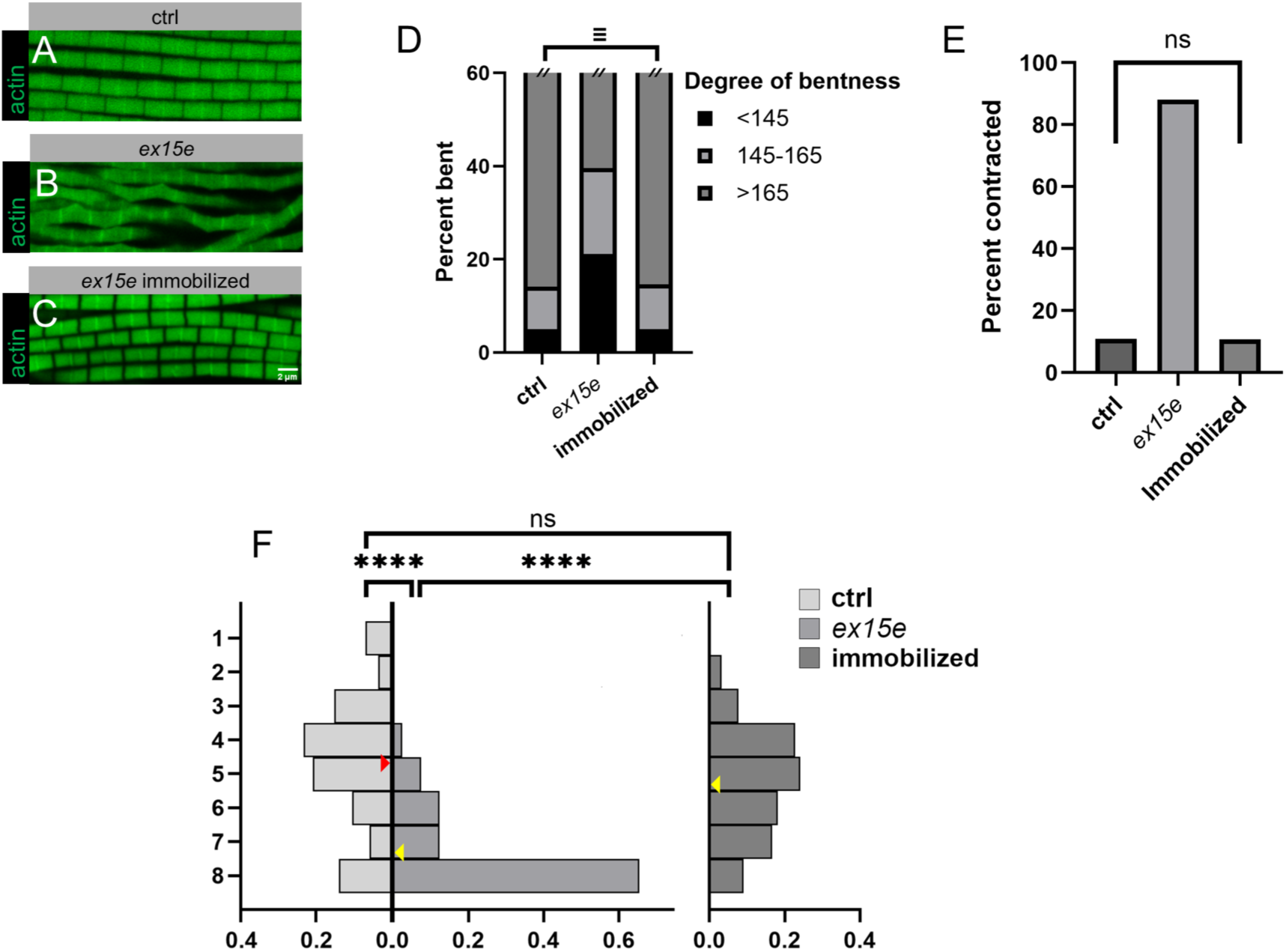
*ex15e* defects are rescued by immobilization. Rescues were attempted by immobilizing flies thereby preventing IFM use. *ex15e* myofibrils in flies that were immobilized **(C)** resembled those of ctrl **(A)**, both showing no phenotypes compared to *ex15e* **(B)**. **(D)** Bend angles were equivalent between ctrl and *ex15e* immobilized flies (*n*=2119) (Two One-Sided Test (TOST) with ±4.5° equivalence margin, α=0.05; *t_L_=*15.36, *t_U_*=-12.00). **(E)** The proportion of sarcomeres that were contracted was not significantly different between ctrl and immobilized flies (*n*=2246) (Fisher’s exact test; *p*=0.9037). **(F)** Flight ability was rescued with immobilized flies (*n*=67 flies) having no significant flight defect compared to ctrl (*n*=86) but being significantly improved from their non-immobilized counterparts (*n*=81) (Fisher’s exact test; ctrl-immobilized *p*=0.0640; ctrl-*ex15e* and *ex15e*-immobilized *p*<0.0001).

## Discussion

Traditionally, IDRs were associated with the concept of amorphousness, finding themselves in transient structures such as condensates and membraneless organelles. As our understanding of disordered proteins has changed, their role in more disparate contexts has become increasingly appreciated, revealing them as modular hubs, flexible scaffolds, and more (Cortese et al., 2008). Furthermore, their role in disease has spurned great interest in their myriad functions. In this study, we demonstrate the role of an exceptionally large disordered region, Zasp52’s exon 15e, in maintaining stability and integrity in a highly rigid and force-bearing structure – the Z-disc.

First, we showed that the expression of exon 15e is spatiotemporally regulated. Being confined to the adult IFM suggests that exon 15e’s function is uniquely required for the rapid oscillatory contractions in indirect flight. Supporting this is the fact that the linker region containing exon 15e is generally much longer in Drosophilids, and rather short in insects using direct flight. Although this linker region is predicted by multiple software programs to be disordered, within it are regions of greater order, one of which corresponds to a putative actin-binding motif (Ashour et al., 2023). This arrangement exists in other proteins such as Cno in Drosophila, where higher-ordered actin-binding regions are embedded in a large IDR (Jensen et al., 2025). Furthermore, a synthetic enhancement between *ex15e* and *Act88F^KM88^* heterozygotes indicates that exon 15e and actin genetically interact. There may be other important short linear motifs (SLiMs) contained within the linker region which serve as interacting sites with other sarcomere proteins. Indeed, multiple sequence alignment of the linker regions of various insects reveals several conserved motifs of unknown function. The presence of an actin-binding motif that may strengthen thin filament anchoring to the Z-disc would be in line with the phenotypes we observed in *ex15e*. However, we were unable to detect defects in our deletion of this motif which supports the theory that the entire IDR is essential and that the ABM is not the key to its function. It is still possible that the ABM within exon 15e acts in concert with the N-terminal ABM, but specific amino acids mediating the N-terminal actin binding have not yet been identified (Liao et al., 2020). Bending at the Z-disc is indicative of a loss of thin filament anchoring and stability at the Z-disc, allowing them to shift out of alignment. Normal sarcomeres bent due to force would exhibit a kinked-hose appearance, where the inner edge is sharply bent and the outer one bowed. This was occasionally observed in controls, whereas *ex15e* bends were always shifted uniformly. Actin in the H-zone is also indicative of loss of stable thin-filament anchoring. During preparation, IFMs are incubated overnight in skinning solution during which they enter a state of rigor. After washing them with an ATP-containing relaxing solution, the myosin heads detach from the thin filaments and the supple nature of the sarcomere which provides it with elastic recoil, causes it to resume a relaxed conformation. However, when thin filaments are weakly anchored at the Z-discs, the natural tension contained within the sarcomere is reduced and thus it will not “bounce back” into a relaxed conformation, resulting in a hypercontracted state which manifests as actin continuing to reside in the H-zone. Cheerio, a Drosophila filamin, was shown to stably attach actin thin filaments at the Z-disc and induces a similar actin incorporation into the H-zone when mutated (González-Morales et al., 2017). Thus, we also demonstrate the importance of Z-disc integrity in allowing sarcomeres to return to their relaxed state in rapidly contracting asynchronous IFM. Accordingly, synchronous muscles do not have such regularly structured Z-discs, and do not require Zasp52’s exon 15e. Indeed, exon 15e is absent in larval and other muscle types. We therefore propose that exon 15e is uniquely required to stabilize rapidly contracting asynchronous IFM.

The orthologous human ALP/Enigma family consists of seven members, with LDB3 (ZASP) considered the canonical Zasp52 ortholog. Zasp52 is more precisely a combined ALP/Enigma member; its first LIM domain is sequentially homologous to the single LIM domain of human ALP family members, and its last three LIM domains are homologous to the three Enigma family LIM domains. Though these human proteins do contain disordered linkers connecting their structured domains, they are much shorter than in Drosophila and are not sequentially homologous. The length of Drosophila’s linker was likely evolutionarily driven to be large, as we see insects using direct flight to have much shorter linkers. Several pathological mutations have been identified in the linker region between LDB3’s PDZ and LIM domains (Vatta et al., 2003), some of which are now known to lie in the actin-binding Zasp motif (ZM) (Lin et al., 2014), which corresponds to the actin-binding region N-terminal to Zasp52 LIM1 (Liao et al., 2020). Interestingly, LDB3 has three skeletal muscle-specific splice isoforms which are differentially expressed and variably retain/omit exons 9 through 11 of a linker sequence (Lin et al., 2014). The prenatal long (L) isoform contains exons 10 and 11, post-natal long (LΔex10) has only exon 11, and short (S) has only 9. Post-natal LDB3-LΔex10 contains an actin-binding region in its linker region whose function is abolished by the inclusion of exon 10 in LDB3-L. Thus, human Zasp protein linkers also function in actin binding, regulated by developmentally differential expression. Linker regions in several human ALP/Enigma proteins were found to interact with both α-actinin and actin using AlphaFold (Healy & Collins, 2023). The mouse ortholog Cypher also contains tissue-specific splice variants which variably include or omit linker sequences (Huang et al., 2003). Like Zasp52’s exon 15e, Cypher is not required for sarcomerogenesis or Z-disc assembly but only maintenance (Zhou et al., 2001).

Our genetic rescues support our hypothesis of exon 15e involvement in stabilizing thin filaments and Z-discs. The rescues also demonstrate that the conserved domains (PDZ, ZM, LIM1-4) are not sufficient for proper Z-disc and thin filament stability in IFM, because Zasp52-PR could not rescue any of the defects. Interestingly, these phenotypes, as well as the flight defects we observed, worsened with age, with defects in one-day-old flies being very minor. This implies that exon 15e is not required developmentally, but rather for the maintenance of sarcomeres which naturally deteriorate with use. We originally hypothesized that since IDRs often lead to protein aggregation and phase separation, exon 15e might drive the formation of Z-bodies, primitive Z-discs which form through the coalescence of diffuse Z-disc proteins including Zasp52. However, given the lack of defects in early Z-discs and the fact that exon 15e expression does not temporally coincide with Z-body formation, we reject this hypothesis. The role of exon 15e in maintenance is supported by immobilization rescues, in which flies were prevented from flying. We opted for a low-ceiling apparatus which we believe would not only prevent flies from flying but discourage them from futilely attempting to utilize their IFM at all. Surprisingly, immobilization restored flight ability to wild type levels. Interestingly, human zaspopathies often manifest late in life, with one study citing onset at 44 to 73 years of age (Selcen & Engel, 2005), much like what we observed in flies. Our data suggest that avoiding strenuous exercise might delay the onset of zaspopathies. Given the maintenance-specific role of exon 15e, but also that as a whole, Zasp52 is indispensable for pupal myofibril development and not just maintenance (Katzemich et al., 2013, Medina-Quintana et al., 2026), Zasp52 appears to have a multifaceted role more complex than previously considered.

It was demonstrated in Liao et al., 2016 that various structured domains of Zasp52, such as the C-terminal LIM domains, are sufficient to localize to the Z-disc. We found that the disordered exon 15e was also able to localize to the Z-disc indicating that some feature contained within it is capable of targeting the protein to its correct location. We took advantage of this to assess its kinetics using FRAP and found that the exon 15e construct is much less dynamic than Zasp52-PR (which lacks exon 15e) suggesting that this disordered region is important for anchoring the protein in place. This finding supports the theory that the IDR acts to stabilize thin filaments at the Z-disc. Although one-day-old *ex15e* myofibrils are morphologically indistinguishable from controls, we were already able to detect abnormalities in Zasp52 dynamics using FRAP in our mutant background. The presence of recovery suggests that certain Zasp52 isoforms are not immobile as might be assumed for an anchoring protein and do indeed possess kinetics on a sub-minute timeframe. This is supported by significant transcription of the gene, including exon 15e-containing isoforms, into adulthood. The mobile fraction of Zasp52-PR was significantly increased in a system entirely lacking exon 15e (the PR isoform also lacks exon 15e) indicating a role of exon 15e in retaining Zasp52 at the Z-disc, whether by direct interaction or not. The FRAP data suggest that Zasp52’s IDR stabilizes the Z-disc, in line with the electron microscopy data. It suggests that Z-discs can be described as highly solid aggregates in part stabilized by IDRs, on the other end of the spectrum of the transient condensates initially described, thereby further expanding the importance of IDRs for establishing subcellular structures.

Overall, Zasp52 and its isoforms represent a complex toolbox of multifunctional proteins. There has been evidence suggesting that Zasp52 is not required for initial sarcomere assembly in embryos (Rui et al., 2010), important for Z-disc maintenance overall (Rui et al., 2010, Jani & Schöck, 2007, Katzemich et al., 2013), and required for pupal myofibril assembly (Katzemich et al., 2013). The complete picture of Zasp52’s varied function is still unclear; however, we have shown that a specific disordered region is required for maintaining IFM and stabilizing the rigidity of Z-discs. The long linker might not only contain protein-interacting regions and provide the glue to hold the Z-disc together but might also provide the spacing needed for the neighboring structured domains to properly interact with their partners and perform their roles. We have seen in our genetic interaction experiment an inability of thin filaments to properly elongate when Act88F is partially depleted, which further implicates exon 15e not only in Act88F structure but dynamics. Most encouraging is the complete rescue of flight ability through immobilization, which may provide avenues for delaying the onset of human zaspopathies through reducing strenuous exercise.

## Materials and Methods

| Key Resources Table |  |  |  |  |
| --- | --- | --- | --- | --- |
| Reagent type (species) or resource | Designation | Source or reference | Identifiers | Additional information |
| gene (*Drosophila melanogaster*) | Zasp52 | FlyBase | FBgn0265991 |  |
| gene (*Drosophila melanogaster*) | Act88F | FlyBase | FBgn0000047 |  |
| strain, strain background (*D. melanogaster*) | *ex15e* | This paper |  | See Material and Methods-Fly strains and husbandry |
| strain, strain background (*D. melanogaster*) | *Zasp52<sup>MI00979</sup>* | BDSC | RRID:BDSC_33099 |  |
| strain, strain background (*D. melanogaster*) | *Df(2R)BSC427* | BDSC | RRID:BDSC_24931 |  |
| strain, strain background (*D. melanogaster*) | $\Delta ABM$ | This paper | | See Material and Methods-Fly strains and husbandry |
| strain, strain background (*D. melanogaster*) | UAS-Zasp52-PF | This paper |  | See Material and Methods-Fly strains and husbandry |
| strain, strain background (*D. melanogaster*) | UAS-Zasp52-PR | This paper |  | See Material and Methods-Fly strains and husbandry |
| strain, strain background (*D. melanogaster*) | UH3-GAL4 | Singh et al., 2014 | FBti0148868 |  |
| strain, strain background (*D. melanogaster*) | *Act88F<sup>KM88</sup>* | FlyBase | FBal0000272 |  |
| strain, strain background (*D. melanogaster*) | UAS-GFPexon15e | This paper |  | See Material and Methods-Fly strains and husbandry |
| strain, strain background (*D. melanogaster*) | UAS-GFPZasp52-PR | González-Morales et al., 2019 | FBtp0140762 |  |
| strain, strain background (*D. melanogaster*) | *w<sup>1118</sup>* | FlyBase | FBal0018186 |  |
| antibody | anti-Zasp52 full-length (Rabbit polyclonal) | Jani & Schöck, 2007 |  | 1:200 |
| antibody | anti-Zasp52LIM234 (Rabbit polyclonal) | This paper |  | See Material and Methods-Western blotting |
| antibody | anti-βPS integrin<br>(Mouse monoclonal) | DSHB | CF.6G11 | 1:200 |
| antibody | Goat anti-rabbit<br>IgG (H+L) HRP | ThermoFisher<br>Scientific | Cat 31460 | 1:10000 |
| antibody | Alexa Fluor 647<br>goat anti-rat | ThermoFisher<br>Scientific | Cat A-21247 | 1:400 |
| antibody | Alexa Fluor 488<br>goat anti-rabbit | ThermoFisher<br>Scientific | Cat A-11008 | 1:400 |
| antibody | Acti-stain 555<br>fluorescent<br>phalloidin | Cytoskeleton<br>Inc | Cat PHDH1 | 1:400 |
| antibody | FluoTag®-X4<br>anti-GFP | NanoTag<br>Biotechnologi<br>es | Cat N0304 | 1:200 |
| software,<br>algorithm | IBS 2.0 | Xie et al.,<br>2022 |  |  |
| software,<br>algorithm | SMART | Letunic &<br>Bork, 2018 |  |  |
| software,<br>algorithm | Salmon 1.10.0 | Patro et al.,<br>2017 |  |  |
| software,<br>algorithm | AIUPred v0.1 | Erdős &<br>Dosztányi,<br>2024 |  |  |
| software,<br>algorithm | DISOPRED3 | Jones &<br>Cozzetto,<br>2015 |  |  |
| software,<br>algorithm | metapredict<br>online v3.0 | Emenecker et<br>al., 2021 |  |  |
| software,<br>algorithm | AlphaFold v2.0 | Jumper et al.,<br>2021 |  |  |
| software,<br>algorithm | iTOL v.6 | Letunic &<br>Bork, 2024 |  |  |
| software,<br>algorithm | NARDINI | Cohan et al.,<br>2022 |  |  |

### Fly strains and husbandry

Flies were housed at room temperature (22°C) in standard glass vials and food unless indicated otherwise. Immobilized flies were reared in between 3.25×3.25 inch glass plates separated by 1.5 mm spacers; standard food was inserted into the gap. Immobilized flies were transferred to a new apparatus with fresh food every couple of days. *ex15e* flies were created by Genetivision using CRISPR-mediated homology-directed repair. The gRNA sequences used were TTTGAACTTGAACTCTAATGTGG for the downstream target and CAGCCTCTCCACCTTTCCGATGG for upstream. The deletion was replaced with a cassette containing a 3xP3DsRed marker flanked by attP landing sites. It was validated by sequencing at the cut sites. *ΔABM* flies were created by Genome ProLab using CRISPR-mediated homology-directed repair as well. The gRNA sequence GAGGAGCTCTCGGGTCTGCA was used to precisely delete five amino acids (LQREL), verified by sequencing. The following genotypes were used: *ex15e*, *ΔABM*, *w^1118^* as control in all experiments, *Df(2R)BSC427*, *Act88F^KM88^*, UAS-GFPZasp52-PR, UH3-Gal4, UAS-Zasp52-PF, UAS-Zasp52-PR, UAS-GFPexon15e, and *Zasp52^MI00979^*. UAS-Zasp52-PF and UAS-Zasp52-PR consist of the respective Zasp52 isoform plus an N-terminal ALFA-tag and C-terminal Spot tag. UAS-GFPexon15e encompasses all amino acids encoded by exon 15e except the last 9 encoding the beginning of LIM2. All UAS constructs were made by Genscript and injected into *w^1118^* by GenomeProlab.

### Western blotting

Extracts were prepared by homogenizing ten flies/fly parts in 100 µL RIPA buffer with protease inhibitor plus an equal amount of 2× Laemmli buffer with β-mercaptoethanol. Samples were boiled at 95°C for five minutes and centrifuged at 12,000×rcf for 5 minutes. Supernatant was used for electrophoresis according to standard protocol using a 4-15% BioRad TGX Stain-free gel. Stain-free gels were imaged for total protein content on a BioRad ChemiDoc before protein was transferred onto a nitrocellulose membrane. Standard Western blotting procedures were followed. Primary antibodies used were rabbit anti-Zasp52 full-length (Jani & Schöck, 2007), and a rabbit antibody against the three C-terminal LIM domains of Zasp52 (FEKYLAPTCSKCAGKIKGDCLNAIGKHFHPECFTCGQCGKIFGNRPFFLEDGNAYCEADWNELFTTKCFACGFPVEAGDRWVEALNHNYHSQCFNCTFCKQNLEGQSFYNKGGRPF CKNHAR) made by Genscript; secondary antibody was goat anti-rabbit IgG (H+L) HRP (ThermoFisher Scientific).

### Locomotory assays

For flight assays, flies were reared in vials (unless immobilized) until the desired age, with vials flipped whenever pupae began to appear. Flight assays were conducted in a manner similar to that described in Babcock & Ganetzky, 2014. A tube 22 cm long and 3.2 cm in diameter was placed in a funnel; the bottom of the tube rested 4 cm above the bottom spout of the funnel. The funnel rested in a 60 cm long tube which was 13.5 cm in diameter and covered with TAD Shake & Spray glue (LADD Research) on the inside. The bottom of the funnel was 12 cm below the top of the larger tube. The larger tube was divided vertically into seven 8.5 cm segments, with segment 1 being at the top, and segment 7 being at the bottom. A dish containing water was placed below the tube and was considered segment 8. Vials containing flies were dropped down the narrow tube so that they would be ejected into the larger tube and land somewhere along its length. Flies naturally fly upwards and those with stronger flight would land higher up the tube. Binary flight assays were conducted by releasing flies one at a time and recording whether they could generate any upwards lift. They were raised in the same conditions as above.

For larval crawling assays, third instar larvae of the desired genotype were collected from bottles and assayed according to Post & Paululat, 2018. In brief, videos of larvae on a slightly moistened petri dish were taken. Time segments where larvae were crawling in a straight line continuously for at least ten seconds were analyzed – contractions per distance were measured and recorded.

### Muscle preparation and immunofluorescence

Primary antibodies used were rabbit anti-Zasp52 full-length (Jani & Schöck, 2007) and mouse anti-βPS-integrin (CF.6G11; obtained from Developmental Studies Hybridoma Bank, Brower et al., 1984) at a concentration of 1:200 for both. Secondary antibodies used were Alexa Fluor 647 goat anti-rat (ThermoFisher Scientific) and Alexa Fluor 488 goat anti-rabbit (ThermoFisher Scientific) at a concentration of 1:400.

For IFM tissue, a protocol similar to Xiao et al., 2017 was followed (refer to for recipes). Briefly, thoraces with bisected and incubated in Relaxing-Glycerol solution at -20°C overnight. IFM were isolated and fixed in 4% paraformaldehyde for 15 minutes. Fixed IFM were incubated in primary antibodies at 4°C overnight. IFM were incubated in Acti-stain 555 fluorescent phalloidin (Cytoskeleton Inc) and secondary antibodies, at room temperature for one hour. Samples were mounted in Mowiol mounting medium.

For thorax hemisections including MTZ, a scalpel was used to sagittally bisect thoraces. Samples were fixed in 4% paraformaldehyde for 15 minutes and incubated in primary antibodies for one hour at room temperature, and then Acti-stain 555 fluorescent phalloidin and secondary antibodies for one hour at room temperature. Samples were mounted in Mowiol mounting medium using three 0.12 mm spacers.

For TEM, bisected thoraces were treated with 5 mM MOPS, 150 mM KCl, 5 mM EGTA, 5 mM MgCl_2_, 5 mM ATP, 1% Triton X-100, and protease inhibitor for two hours. then the same solution with 50% glycerol instead of Triton X-100 overnight at 4°C. This was repeated at -20°C. Samples were washed in 5 mM MOPS, 40 mM KCl, 5 mM EGTA, 5 mM MgCl_2_, 5 mM NaN_3_, protease inhibitor, and then fixed in 3% glutaraldehyde and 0.2% tannic acid for two hours. Samples were washed in 20 mM MOPS, 5 mM EGTA, 5 mM MgCl_2_, 5 mM NaN_3_, and then in 100 mM sodium phosphate buffer, 5 mM MgCl_2_, 5 mM NaN_3_. Secondary fixation was performed in 1% osmium tetroxide in the previous sodium phosphate buffer for one hour. Samples were stained in 2% uranyl acetate, washed with water, and underwent a serial dehydration in ethanol. Samples were washed with propylene oxide and serially infiltrated with Mollenhauer Epon-Araldite. After curing in Beem capsules, 70 nm sections were cut and stained in 4% uranyl acetate followed by SATO lead stain.

### Fluorescence Recovery After Photobleaching

The sequence encoding exon 15e minus the nine C-terminal amino acids encoding LIM2 was subcloned into pUAST-attB-GFP-V5-His and injected into *w^1118^*. IFM were prepared as previously described (with fixation) and stained with ActiStain-555 and FluoTag^®^-X4 anti-GFP (NanoTag Biotechnologies) to visualize localization; FRAP was performed as described below. One-day-old female flies were anaesthetized and their abdomens, wings, and legs removed. Carcasses were pinned down by the head on a silicone dish and immersed in hemolymph-like 3 (HL3) saline (Stewart et al., 1994). Thoraces were sagittally bisected, washed thrice in HL3 saline, and mounted with a 0.12 mm spacer. Imaging was performed immediately using a HC PL APO 63×/1.40 OIL CS2 objective on a Leica SP8 point-scanning confocal microscope. Five pre-bleach frames (1.29 seconds each) were recorded before bleaching with a 488-nm laser to achieve a bleach depth averaging 75%±4% (SD) for UH3>GFPexon15e, 74%±8% for UH3>GFPZasp52-PR, 77%±7% for *ex15e*; UH3>GFPZasp52-PR, and 76%±9% for fixed samples. Seventy-five post-bleach frames (1.29 seconds each) were then acquired. ROIs (1.25×3 µm) were placed over individual Z-discs and those with drift were discarded. For each fly, three Z-discs on different myofibrils were bleached, and intensities of adjacent unbleached Z-discs and unbleached background regions were recorded. Fluorescence was normalized as (bleached–background)/(unbleached–background), averaged across the three ROIs, and scaled such that the post-bleach frame equaled 0 and the pre-bleach mean equaled 100. Recovery curves were presented as mean±SEM. Each fly’s curve was fitted to a one-phase exponential association model to obtain the plateau (mobile fraction). Fixed UH3>GFPZasp52-PR samples were prepared in the same manner except with an additional ten-minute fixation in 4% paraformaldehyde in HL3 saline.

### Microscopy

Confocal images were acquired using a HC PL APO 63×/1.40 OIL CS2 objective on a Leica SP8 point-scanning confocal microscope. The position along the z-axis was adjusted until perfectly in the centre of the myofibril, i.e., the widest point. Polarized light images were acquired using a 5X N-ACHROPLAN, NA=0.13 objective on a Zeiss Axio Observer microscope. TEM images were acquired using an FEI Tecnai G2 Spirit Twin 120 kV Cryo-TEM using an AMT NanoSprint15 MK2 CMOS camera.

### Computational analyses

Schematics were illustrated with IBS 2.0 (Xie et al., 2022). All protein domains were defined per SMART (Letunic & Bork, 2018). Graphing and statistical analysis was performed using GraphPad Prism 10. Statistical analyses were performed in GraphPad Prism 10, where ns=*p*≥0.05, *=*p*<0.05; **=*p*<0.01, and ****=*p*<0.0001, and calculated to four decimal places. Equivalence of bend angles was assessed using the Two One-Sided Test (TOST) procedure. Model sarcomeres with bend angles ranging from 180° (straight) to 95° and blurring were generated. Each was hand measured ten times and the average absolute measurement error (4.5°) was used as the equivalence margin for all TOST analyses.

Minus strand RNA-seq bigWig files from the ENCODE project (Duff et al., 2015) with the following accessions were retrieved and loaded in IGV 2.19.4 (Robinson et al., 2011): ENCFF289ZZT, ENCFF982OGI, ENCFF430HWM, ENCFF716DNU, ENCFF829OGQ, ENCFF936YJM, ENCFF987YVI, ENCFF033MIY, ENCFF845HII, ENCFF021VXY, ENCFF792FNT. Signal tracks were aligned to the dm3 assembly and the data range was adjusted uniformly for all tracks.

SRA files with the following accession numbers were downloaded from the Bioproject PRJNA419412 (Spletter et al., 2018): SRR6314253, SRR6314256, SRR6314275, SRR6314259, SRR6314277, SRR6314262, SRR6314273. Salmon 1.10.0 (Patro et al., 2017) was used to quantify expression of the following transcripts which correspond to identified Zasp52 isoforms: FBtr0392902, FBtr0329914, FBtr0329912, FBtr0100387, FBtr0111048, FBtr0087315, FBtr0329913, FBtr0310086, FBtr0329916, FBtr0329915, FBtr0310088, FBtr0310087, FBtr0392901, FBtr0329911, FBtr0342711, FBtr0100388, FBtr0302163, FBtr0301313, FBtr0479884, FBtr0329909, FBtr0392918, FBtr0329910. The first six were classified as containing only exon 15a, the next six containing exons 15b-d, the next four containing exon 15e, and the last six containing no exon 15.

ImageJ was used to evaluate bending manually using the angle tool, and sarcomere length using the line measuring tool.

For disorder prediction, the entire Zasp52-PF sequence was entered into AIUPred v0.1 (Erdős & Dosztányi, 2024), DISOPRED3 (Jones & Cozzetto, 2015) (as two sequences, the first covering the first 1200 amino acids and the second, the remaining), or metapredict online v3.0 (Emenecker et al., 2021). Raw values for each residue were obtained and smoothed using a Savitzky-Golay filter with fourth order smoothing and 11 neighbours on each side. AlphaFold v2.0 (Jumper et al., 2021) was used to obtain pLDDT scores for the alpha carbon of each residue. pLDDT scores were min-max normalized and smoothed in the same manner as above.

Zasp52 orthologs were obtained for members of the class *Insecta* using NCBI’s Eukaryotic Genome Annotation Pipeline. Canonical protein sequences for each were downloaded and those which did not contain the classical PDZ and four LIM domains were removed. The number of residues between the first and second LIM domains was calculated. A phylogenetic tree of these species was created using NCBI’s Common Tree (Schoch et al., 2020) and a heatmap created using iTOL v.6 (Letunic & Bork, 2024).

NARDINI (Cohan et al., 2022) was used for the sequences indicated to generate a z-score matrix, with 100,000 scrambled sequence variants.

## Acknowledgments

This work was funded by the Canadian Institutes of Health Research project grant PJT-178114. Confocal images were collected in the McGill University Advanced BioImaging Facility (ABIF). Electron microscopy images were collected and analyzed in the McGill University Facility for Electron Microscopy Research (FEMR).

## Data availability

All data used in this study is available on FigShare at https://figshare.com/projects/Supplementary_material_for_Zasp52_s_differentially_expressed_intrinsicall y_disordered_region_confers_thin_filament_stability_at_the_Z-disc/278459.

---

**Figure 1 - Figure supplement 1.**
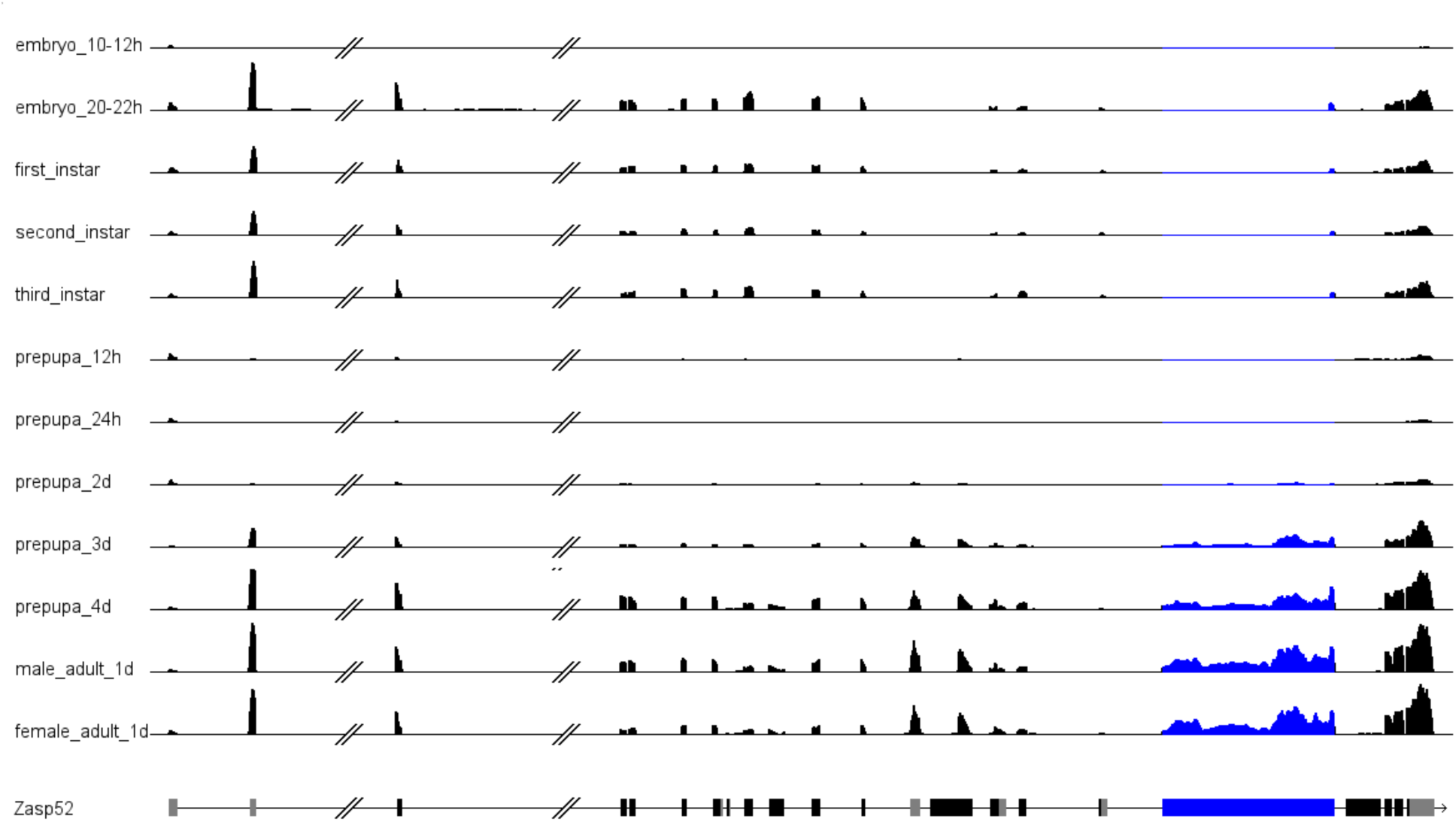
RNA-seq data of exon 15e expression. MODENCODE RNA-seq data showing expression levels at the *Zasp52* locus across various developmental stages. Exon 15e is highlighted in blue.

**Figure 1 - Figure supplement 2.**
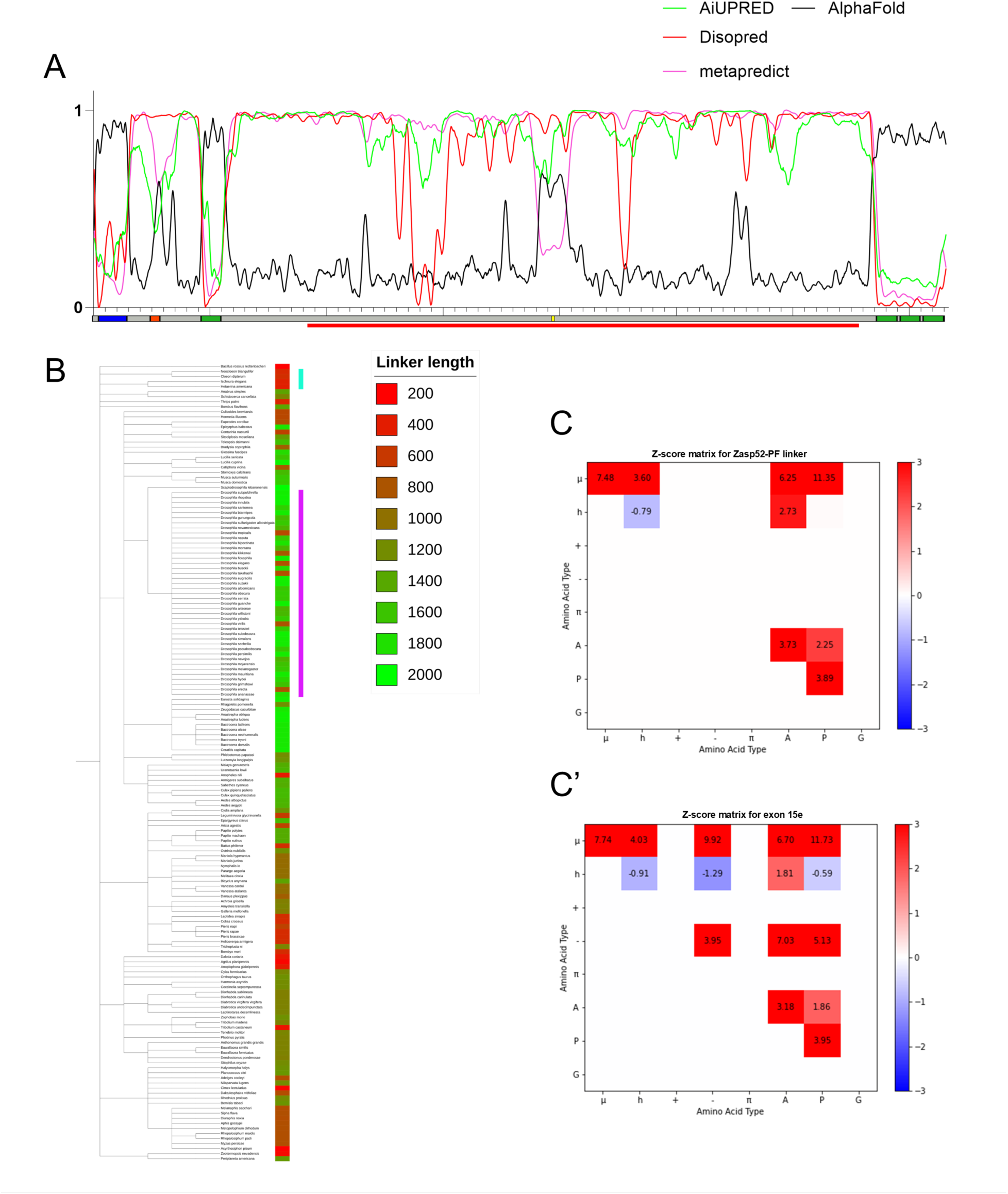
Disorder prediction of amino acids encoded by exon 15e. **(A)** Various disorder prediction algorithms were used to assess the level of per-residue disorder at the *Zasp52* locus. The corresponding *Zasp52* locus is schematized below, with PDZ domain (blue), ZM (red), LIM domains (green), and the ABM identified in Ashour et al., 2023 (yellow). The red bar below the Zasp52 schematic indicates the extent of the deletion in *ex15e*. For disorder prediction algorithms (AiUPRED, Disopred, and metapredict), the higher y-values represent higher levels of disorder while low values close to zero represent low levels of disorder (structure). AlphaFold pLDDT scores are plotted alongside in black. pLDDT scores can be used to inform about disorder with higher scores indicating higher confidence that a region is ordered, and lower scores suggesting disorder (Ruff & Pappu, 2021; Emenecker et al., 2021). **(B)** Dendrogram of various insects with Zasp52 orthologs that contain a PDZ domain and four LIM domains. Canonical isoforms which usually are also the longest ones were used to compute the number of residues comprising the linker between the first and second LIM domains, with data represented in a heatmap. The purple bar indicates all Drosophilids, and the teal bar indicates species of the order *Ephemeroptera* and *Odonata* which use direct flight. Though some Drosophilids contain shorter linkers, no other species contain linkers of comparable length to Drosophilids and other closely related Dipterans. NARDINI Z-score matrices of the linker region between LIM1 and LIM2 in Zasp52-PF **(C)** and of only the sequence encoded by exon 15e **(C’)**. The diagonal squares represent Ω-values which quantify the linear mixing versus segregation of all of residue type *x* with respect to all other residues. The off-diagonal squares represent δ-values which quantify the linear mixing versus segregation of pairs of residue types. A high z-score thus indicates segregation/blocking of a certain residue type with respect to itself/others compared to a null model; a low score indicates even dispersion and intermixing. Polar residues µ = {S,T,N,Q,C,H}; hydrophobic residues h = {I,L,V,M}; basic residues + = {R,K}; acidic residues - = {E,D}; aromatic residues π = {F,W,Y}; alanine = A; proline = P; glycine = G.

**Figure 2 – Figure supplement 1.**
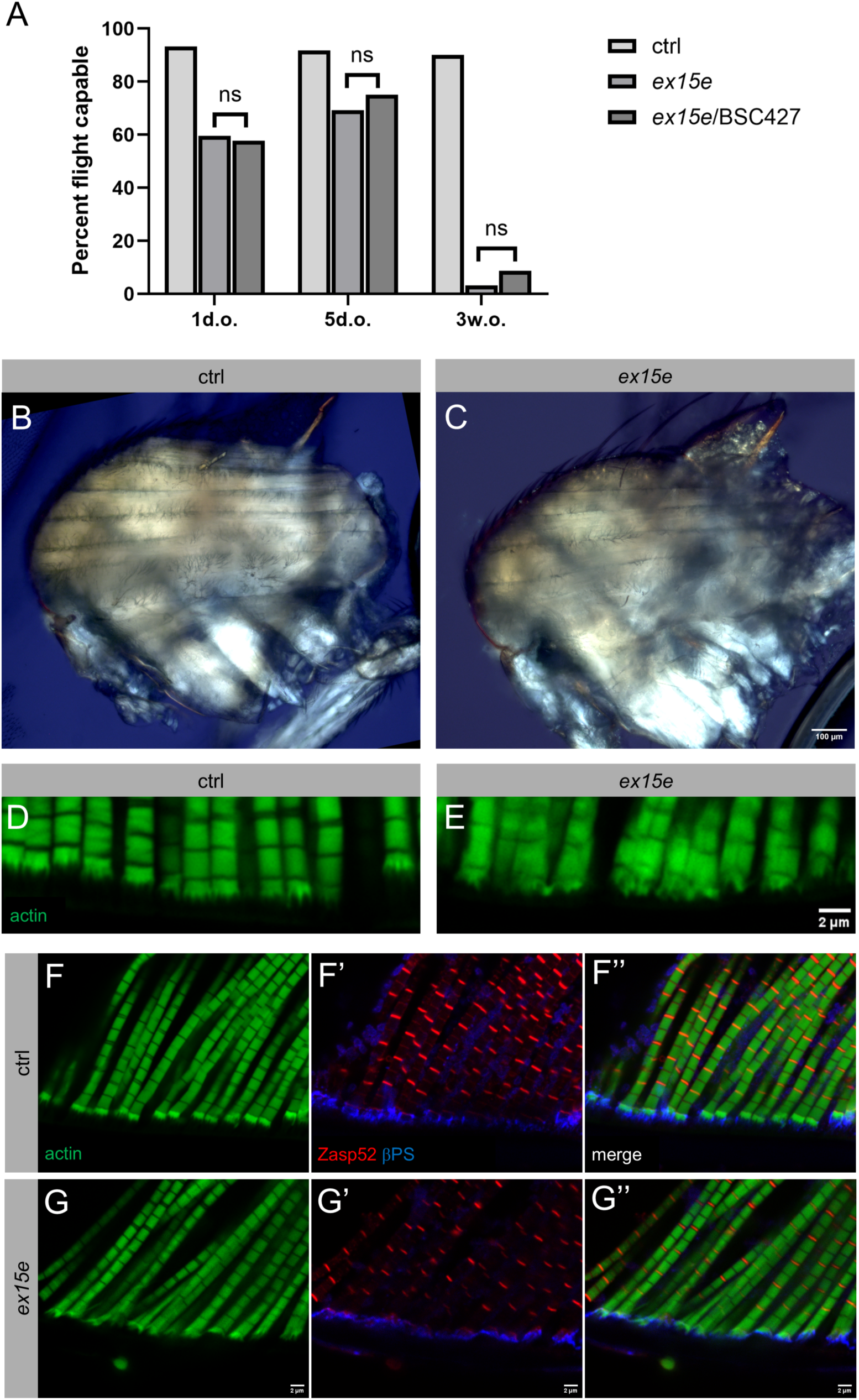
Flight assays, gross muscle morphology and modified terminal Z-discs in *ex15e* mutant. **(A)** A binary flight assay across three age categories of ctrl, homozygous *ex15e*, and *ex15e* over the Zasp52 deficiency *BSC427*. Sample size were as follows: one-day-old (ctrl *n*=44 flies, *ex15e n*=42, *ex15e*/*BSC427 n*=45), five-days-old (ctrl *n*=48, *ex15e n*=39, *ex15e*/*BSC427 n*=28), and three-weeks-old (ctrl *n*=61, *ex15e n*=30, *ex15e*/*BSC427 n*=23). Flight was scored manually on whether they could generate upwards lift or not. There was no significant difference between the homozygous mutant or *ex15e* over the deficiency (Fisher’s exact test; 1 d.o. *p*=1.0000; 5 d.o. *p*=0.7843; 3 w.o. *p*=0.5729). Polarized light images of sagittally bisected thoraces of three-week-old ctrl **(B)** and *ex15e* **(C)** revealed no obvious differences in gross morphology; females shown. **(D-G’’)** Modified terminal Z-discs (MTZ) were affected in *ex15e*. Close up views revealed a disruption in the MTZ and adjacent myofibrils in three-week-old *ex15e* mutants **(E)** compared to ctrl **(D)**. The regular “crown”-shaped structure of the ctrl MTZ is replaced with a disordered structure of variable shape in *ex15e* mutants. βPS integrin and Zasp52 localization appeared unaffected in *ex15e* **(F-G’’)**. Thin filaments were visualized with phalloidin in green, Z-discs with a Zasp52 full-length antibody in red, and integrin adhesion sites at MTZs with a βPS integrin antibody in blue.

## Notes

### Competing Interest Statement

The authors have declared no competing interest.

### Summary of Updates

new figure 5, additional data for figure 8, other modifications

